# Dynamics and geometry of emotion and cognition: an interpretable model of individual human behavior

**DOI:** 10.64898/2026.09.08.749516

**Authors:** Jinyung Hong, Elliot M. Nester, Lekha Varisa, Zuzanna Wrobel, Kris Phataraphruk Rains, Tejas Umesh, Tamera R. Schneider, Sarah H. Lisanby, Nicole A. Roberts, Pavan Turaga, Gene A. Brewer, Andrew I. Yang

## Abstract

Affective flexibility–the capacity to flexibly process emotional information–has been assessed with affective task switching, in which response times consistently reveal larger behavioral costs when switching to the affective task. These traditional metrics of task behavior, however, discard the sub-trial dynamics that produce them and are vulnerable to trial-sampling noise. Subject-specific generative models were independently fitted (22 adults) with response-time sequences. The learned dynamics were visualized as continuous trajectories in a labeled embedding that was shared across person-specific models, yielding a suite of interpretable geometric metrics that captured how sub-trial behavior is jointly shaped by experimental conditions and between-subject variability. Switch–repeat onset distance tracked individual differences in switch costs; early- and late-epoch tortuosity provided convergence with classical sub-process models; and angular alignment between trajectories supported a task set flexibility account of asymmetric switch costs. Overall, our approach demonstrates the utility of latent geometry for deriving mechanistically informative measures of individual task behavior.

## 1 Introduction

Affective flexibility (AF) is the ability to adaptively engage with, or disengage from, the emotional aspects of a stimulus or situation for goal-directed behavior. It is a capacity lying at the intersection of cognitive flexibility and affective processing, and is of higher ecological relevance compared to each domain in isolation. An established behavioral paradigm to test AF is affective task switching (ATS), whereby subjects flexibly switch between an affective task and a neutral task, commonly implemented with subjects viewing images of naturalistic faces (targets) while switching between cue-directed judgments of the face’s emotional expression or gender ^(21;20;48)^. As in other task switching paradigms, switching incurs a behavioral cost–switch cost– related to task set reconfiguration. Asymmetry in switch costs–with greater costs of switching to the emotional task vs. switching to the gender task–is a robust group-level behavioral hallmark that has been consistently observed in multiple independent cohorts ^(66;21;20;48)^. Although asymmetric switch costs have been observed in neutral task switching (NTS) paradigms as well ^(3;2)^, the underlying neurobehavioral processes are thought to be distinct. In NTS, switching to the more “dominant task” engenders a higher switch cost, where dominance of a task set is assigned based on faster performance when repeating the task and/or less interference from the other task set ^(3;2;77;79)^. In ATS, however, neither factor applies to the task set associated with higher switch costs (emotion task) ^(71;48;61;20;21)^, underscoring that ATS uniquely taxes task set reconfiguration for adaptive processing of stimuli whose affective salience may be goal-relevant or irrelevant.

Switch costs are commonly operationalized by the difference of trial-averaged response times (RTs) between switch trials and the corresponding repeat trials. Sequential sampling models (e.g., the drift–diffusion model (DDM)) have been used to infer sub-trial processes from condition-wise RT distributions ^(62;64)^. DDM decomposes sub-trial behavior into evidence accumulation or non-decision processes, the latter of which includes stimulus perception, motor execution, and, in the case of switch trials, task-set reconfiguration ^(70)^. While DDM parameters can aid in characterizing group-level behavioral effects (e.g., asymmetric switch costs), they have similarly been limited in translation to the study of individual differences ^(35;43)^. A general limitation of these metrics is that they reduce task behavior into coarse condition-wise summaries, and fail to capture the rich temporal dynamics of behavior as it evolves across time from stimulus onset to behavioral response, nor the multiscale dependencies in behavior both within and across trials.

Recent advances in deep learning, artificial neural networks (ANNs), and latent dynamical systems have created new opportunities to model the multiscale temporal dynamics of behavior (and neural activity). ANN models are distinguished from other related approaches–e.g. Bayesian/probabilistic ^(53)^, symbolic/cognitive architecture ^(9)^, connectionist ^(13)^–in that they require fewer structural assumptions and fewer handcrafted components that are subject to bias and subjectivity ^(73)^. However, the majority of ANN applications have followed a task-optimization (goal-driven) paradigm, where models are trained to perform the task itself independently of behavioral data. There is a growing body of work in which descriptive ANN models are fitted directly to real behavioral data to recapitulate realistic human behavior; however, the majority rely on data pooled across subjects, either exclusively ^(16;22;8;50)^ or as group-trained components from which subject-level models are derived ^(40;15;29;36)^. Even in the latter case, subject-level estimates are regularized toward the population structure, attenuating subtle individual differences in behavioral variability, idiosyncrasies, and biases.

Here we introduce GeoDynFormer, a generative deep-learning framework to model subject-specific behavior at both sub-trial and cross-trial (task condition) levels. GeoDynFormer models task behavior as a temporally-evolving latent dynamical process that generates sequences of behavioral responses. Subject-specific models trained on the observed trial-by-trial RTs not only reproduced each subject’s RT distributions across conditions, but also estimated sub-trial behavioral dynamics that are not directly observable in the raw behavioral data. A key feature of GeoDynFormer is that model-generated latent dynamics could be visualized in a lower-dimensional embedding as temporal trajectories (state-space perspective), providing an interpretable scaffold to study how task behavior is jointly shaped by task conditions and sub-optimalities specific to each individual subject. These behavioral patterns were quantified with a suite of intuitive geometric features of latent trajectories.

We applied this framework to ATS behavior to test the hypothesis that both group-level behavioral effects and individual differences can be captured via the geometry of latent trajectories. Latent dynamics mapped onto a three-dimensional, interpretable space whose axes corresponded to ATS conditions in a way that generalized across the independently trained subject-specific models. We demonstrate the utility of GeoDynFormer as a novel measurement approach for task behavior: geometric metrics had superior reliability in capturing individual differences vs. raw switch costs; subtrial latent dynamics revealed the geometric substrates of behavioral sub-processes represented by DDM parameters; the geometry of cross-task set interference provided support for a theoretical account on the origins of asymmetric switch costs that is based on differential task set flexibility when switching between affective and non-affective tasks ^(71)^.

## 2 Results

### 2.1 Latent-state behavioral modeling of affective task switching

We model behavior in the ATS paradigm as a problem of sequential latent-state inference, in which continuous streams of task stimuli–comprised of cues encoding information on task type and targets consisting of naturalistic face images–are mapped onto time-varying behavioral responses (Fig. 1). Rather than treating trials as independent events, GeoDynFormer models behavior with a generative dynamical system that transforms stimulus sequences into response sequences. This formulation allows the model not only to fit subject-specific behavior at the single-trial level, but also to characterize how internal task representations evolve over time within trials.

**Fig. 1:**
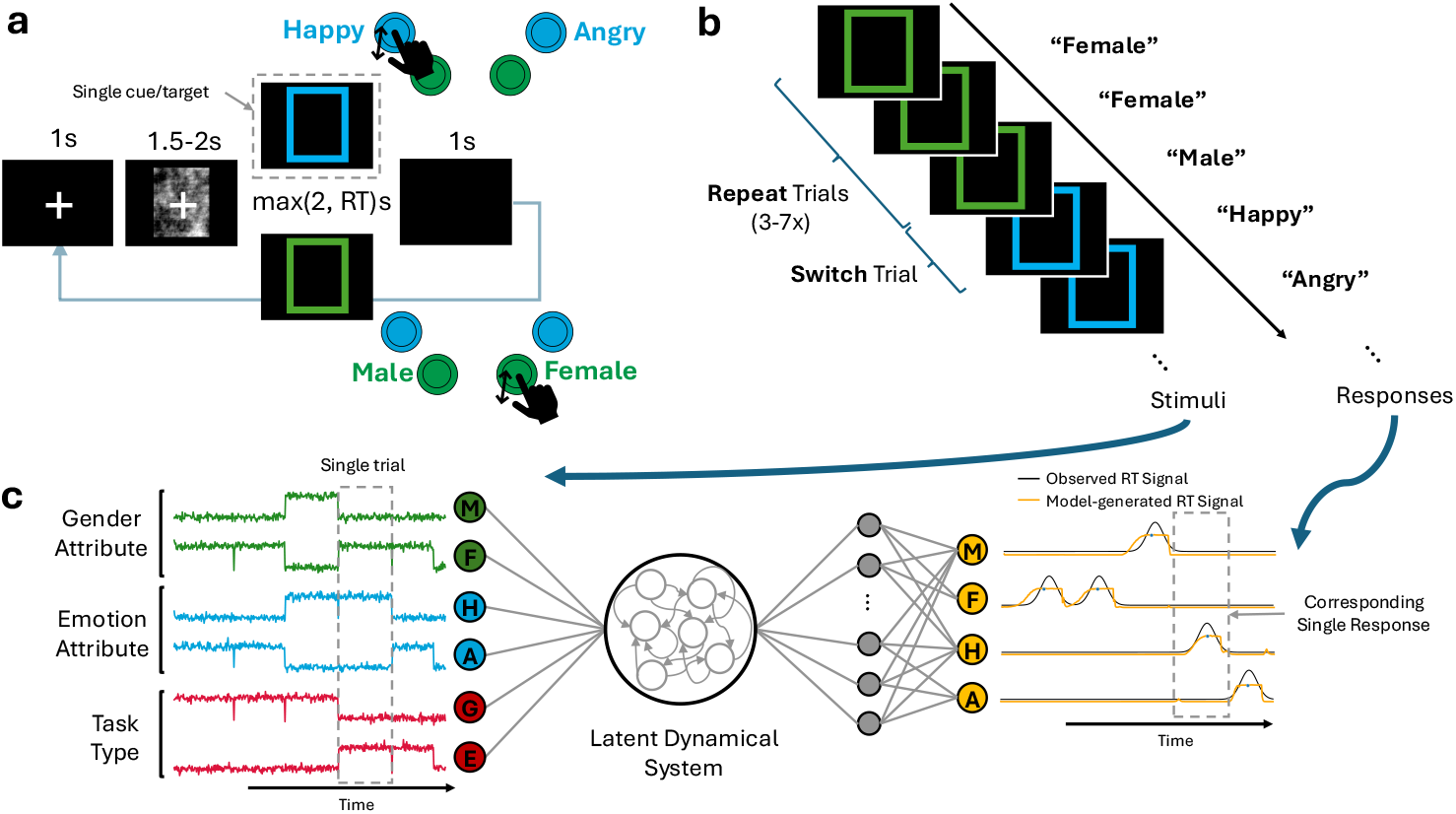
Affective task-switching (ATS) paradigm and modeling framework, GeoDynFormer. **a**, Single-trial structure of the ATS paradigm. Each trial comprised fixation, a scrambled-face interval, and a face target with a colored task cue. Subjects classify the face by emotion (happy/angry) or gender (male/female), followed by a post-response interval. **b**, Example sequence showing several repeat trials followed by a switch trial. **c**, Overview of GeoDynFormer. Left, the converted task cues and face attributes as input streams. Individual trials were input as a multivariate time series with time spanning cue/target onset to RT (grey dashed box corresponding to in (a); see Methods). Right, the signal constructed from the subject’s actual reaction time (black) and the model-generated response signal (yellow). Beyond information about the presented task cue and gender/emotion attributes, model training used only stimulus onset time and observed reaction time, without explicit switch/repeat labels or sub-trial process annotations.

At the core of GeoDynFormer is a latent dynamical system,

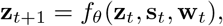

where **z**_*t*_ denotes the latent state, **s**_*t*_ the input stimulus, and **w**_*t*_ a stochastic latent perturbation. As in prior latent dynamical formulations ^(39)^, this state-space framework can capture complex temporal dependencies in task behavior. Notably, GeoDynFormer implements posterior inference over the latent dynamics using Transformer-based attention ^(75)^, rather than the recurrent inference architecture of Jaffe et al. ^(39)^; a matched comparison using the same ATS training framework showed more accurate condition-dependent RT recovery with Transformer-based inference (Extended Data Fig. 1; Methods).

For model fitting, task stimuli were converted into time-varying signal representations. Each task stimuli, both cues and targets (Fig. 1b), were encoded as a dynamic input channel, with a binary-valued unit indicating its presence or absence at each moment in time (Fig. 1c, left). In this representation, information on whether the current trial requires switching vs. repeating the previous trial’s task type was not explicitly provided to the model and therefore had to be inferred from the history of trial sequences. To facilitate training and increase biological plausibility, we added independent zero-mean Gaussian noise (0.1 s.d.) to each stimulus unit.

Model outputs **r**_*t*_ were generated by a decoder distribution *p*_*θ*_(**r**_*t*_ | **z**_*t*_) parameterized by a multilayer perceptron (MLP). Each possible task response was represented by a separate output channel, whose activation at each time point was related to the probability of producing the corresponding response (Methods). To compare model outputs with subject behavior, we transformed each observed RT into a smooth response signal by centering a Gaussian kernel (s.d. = 300 ms) at the subject’s RT, thereby forming a continuous response template (Fig. 1c, right). Parameters of the generative model were then learned using an approximate inference framework based on the variational autoencoder ^(30;67)^ (Methods).

The resulting generative model obeys standard state-space assumptions: given **s**_*t*_ and **w**_*t*_, the current response **r**_*t*_ and future state **z**_*t*+1_ depend only on the current latent state **z**_*t*_. This state-space formulation is important for interpretability because it allows the progression of within-trial computation to be examined directly through the latent state at each moment in time.

We fit an independent GeoDynFormer model to each subject’s behavior on ATS. The training set consisted of each subject’s behavioral stream segmented into sequences of 5 s (corresponding to 2–4 trials), defining the temporal context over which Transformer-based posterior inference operated during training (Methods). After training, each fitted model was used to generate responses on longer, never-before-seen trial sequences from the held-out test set (10 s; Methods), allowing comparison of the generated responses with the subject’s observed behavior on the same held-out trial sequences (Fig. 2; Extended Data Figs. 2, 3).

**Fig. 2:**
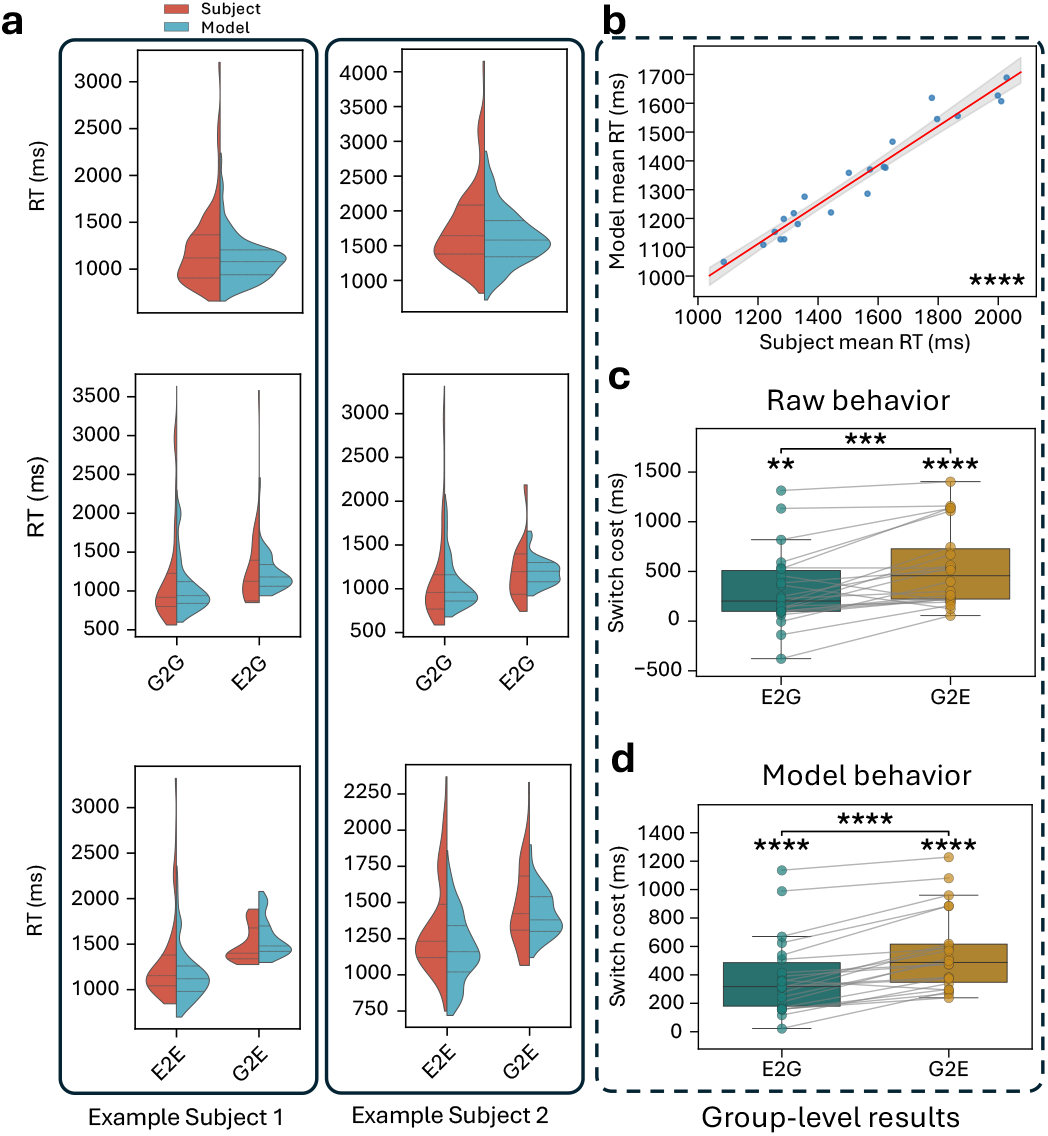
GeoDynFormer captures individual human behavior in ATS. **a**, Observed and model-generated RT distributions for two example subjects, shown for all correct trials (top) and for each condition. **b**, Observed vs. model-generated trial-averaged RTs. Circles indicate individual subjects and their corresponding fitted models; red line, best linear fit; gray shaded region, 95% bootstrapped CI. **c,d**, Observed (**c**) and model-generated (**d**) switch costs for E2G and G2E. Circles indicate individual subjects or models, and gray lines connect paired observations. Box plots show 25th percentile, median, and 75th percentile. One-sample *t*-tests against zero. Horizontal lines indicate paired-sample *t*-tests. ∗∗ *p <* .01, ∗∗∗ *p <* .001, ∗∗∗∗ *p <* .0001.

We first confirmed that GeoDynFormer could reproduce subject-specific task behavior. Subject-level models closely matched the observed condition-wise RT distributions (Fig. 2a). Model-generated and observed grand-average RTs were strongly correlated across subjects (Fig. 2b; *N* = 22, Pearson’s *r* = 0.97, bootstrap 95% CI (0.95, 0.99), best-fit slope = 0.68, *p <* 10^−13^), and similarly for condition-wise RTs (Extended Data Fig. 3a).

GeoDynFormer also reproduced the hallmark asymmetry in switch costs. In the study cohort, switching to the emotion task (G2E) incurred a larger cost than switching to the gender task (E2G) (Fig. 2c; Extended Data Fig. 2). Switch cost is traditionally quantified with the difference between switch-trial and corresponding repeat-trial RTs: *RT*_*E*2*G*_ − *RT*_*G*2*G*_ and *RT*_*G*2*E*_ − *RT*_*E*2*E*_. Across subjects, mean switch cost was 318 ± 84 ms (mean ± s.e.m.) for E2G and 539 ± 87 ms for G2E (paired *t*(21) = −4.26, *p <* 10^−3^). Model-generated responses reproduced this asymmetry (Fig. 2d; Extended Data Fig. 3b), with switch costs of 383 ± 59 ms for E2G and 556 ± 60 ms for G2E (*t*(21) = −7.09, *p <* 10^−6^).

### 2.2 Visualization of latent dynamics in an interpretable lower-dimensional embedding

Having established that GeoDynFormer reproduces subject-specific behavior, we next sought to examine the multiscale organization of its latent states. The sub-trial evolution of latent representations can be represented as continuous trajectories, averaging across repeated trials for each task condition of interest.

We first focused on visualization of latent dynamics in a lower-dimensional embedding. We chose to depict these trajectories in a three-dimensional (3D) uniform manifold approximation and projection (UMAP) embedding ^(54)^; compared to linear methods such as PCA, UMAP accommodates nonlinear structure in the learned latent trajectories, and prioritizes the preservation of local neighborhood relations among latent states ^(34)^. We used the UMAP embedding for visualization and to motivate hypothesis-guided tests of latent-state geometry; unless otherwise noted, all reported statistical tests were performed in the native 16D latent space (Methods).

We depicted trial-averaged trajectories stratified across the two repeat conditions (E2E, G2G) and, within each repeat condition, by the corresponding task-relevant target attribute (male vs. female in G2G; happy vs. angry in E2E). Repeat-condition trajectories were clearly organized along the three ATS-related variables (task type, task-relevant target emotion, and task-relevant target gender); this organization emerged consistently across the UMAP embeddings of independently-trained subject-level models (subject examples in Fig. 3a, Extended Data Fig. 4b).

**Fig. 3:**
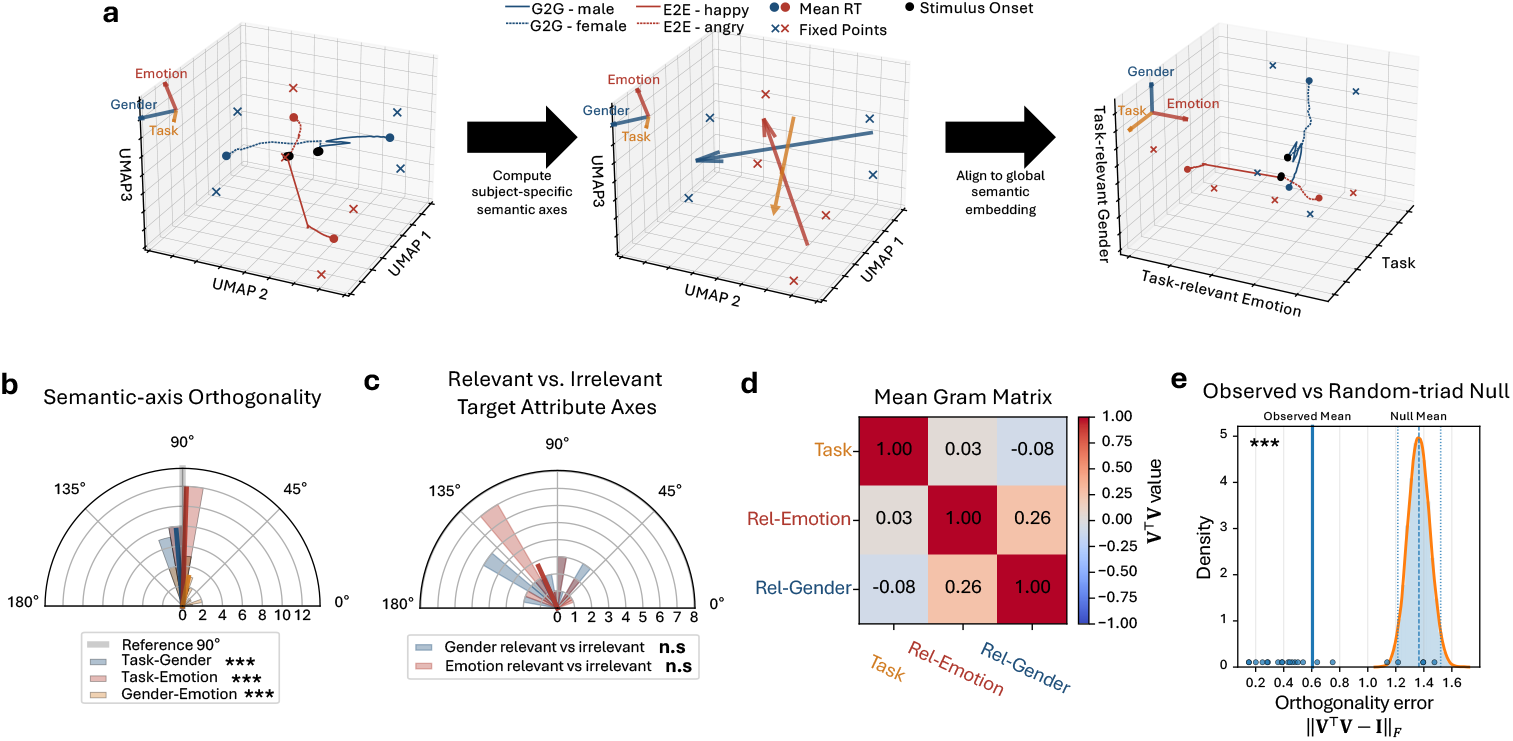
Organization of latent trajectories across semantic axes. **a**, Trial-averaged latent trajectories generated by an individual subject’s model are depicted across repeat task conditions (E2E, G2G) in 3D UMAP embedding (left panel). Each repeat condition is further stratified based on task-relevant target attributes (happy vs. angry for E2E; female vs. male for G2G). Three semantic axes are defined from geometric anchors (see Results; middle panel): Task axis, and task-relevant emotion and gender axes. The subject-specific UMAP embedding is aligned to a common semantic embedding spanned by these three axes (right panel). **b**, Group-level angular relations among the raw task, task-relevant emotion, and task-relevant gender semantic contrast directions, showing approximate orthogonality in 3D UMAP embedding (V-test against 90^◦^). **c**, Group-level angular relations between task-relevant and task-irrelevant emotion and gender contrast directions, showing weak alignment across task types (V-test against 0^◦^). **d**, Mean Gram matrix of the unit-normalized raw semantic contrast vectors in the 3D UMAP embedding. **e**, Subject-level orthogonality error of the raw semantic contrast triad relative to a 3D random-triad null distribution: group-average error = 0.61, null mean = 1.37; 95% null interval (1.22, 1.52). n.s.: not significant, ∗∗∗ *p <* .001.

We leveraged this shared configuration to enhance the interpretability of the embedded dynamics by aligning each subject’s UMAP embedding into a common reference frame with labeled dimensions. To do so, we extracted from each subject’s UMAP space the axes for the three ATS variables–referred to as semantic axes– by computing contrast vectors between the stable fixed points associated with repeat trajectories (Fig. 3a, Fig. 4b; Methods): the task-type axis contrasted the fixed points of G2G and E2E conditions; the task-relevant emotion and gender axes contrasted the fixed points of E2E and G2G repeat trajectories stratified by the corresponding target attributes (e.g., task-relevant emotion axis was anchored by the fixed points of E2E happy and E2E angry trials).

**Fig. 4:**
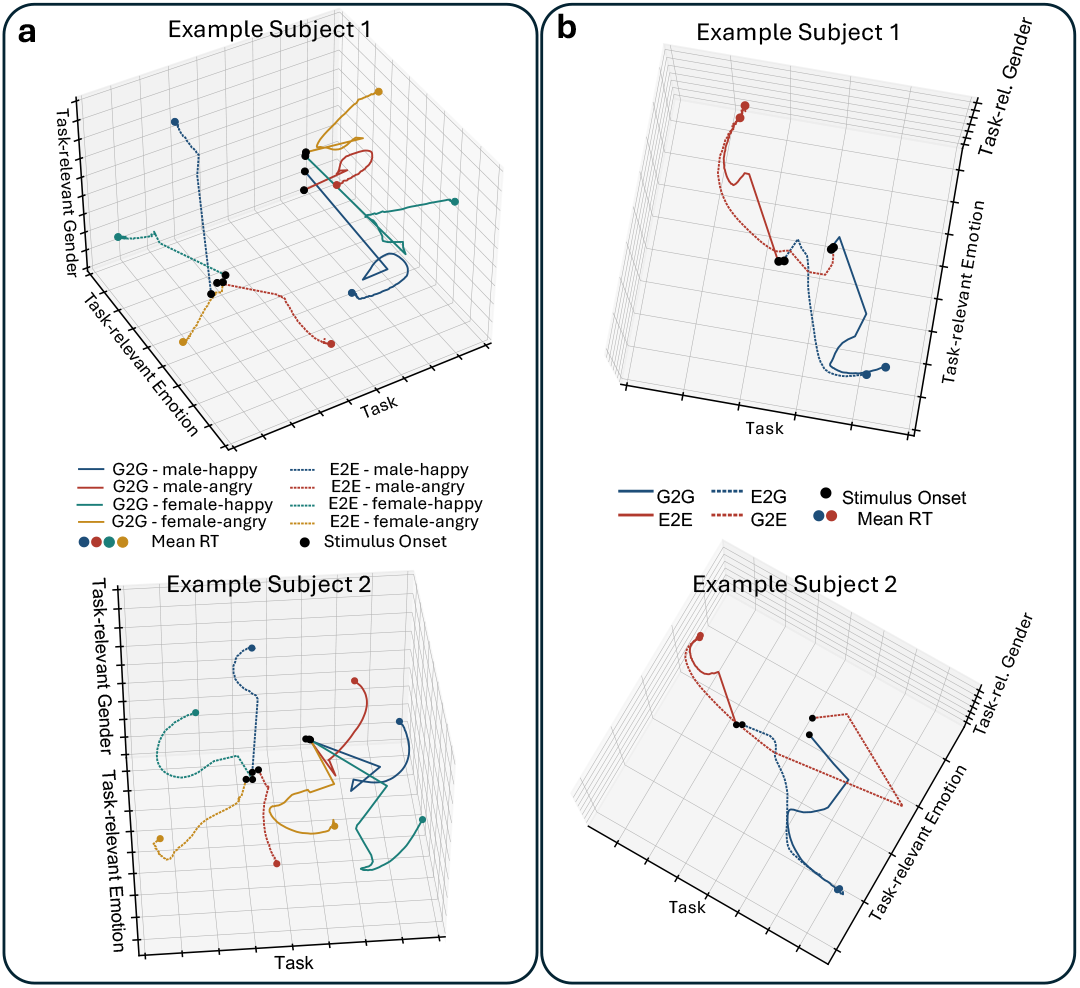
Repeat and switch latent trajectories show interpretable dynamics in common semantic embedding. **a**, Trial-averaged latent trajectories for each repeat condition (E2E, G2G), further stratified across all 4 target attribute conditions (male/female × angry/happy). **b**, Trajectories shown across repeat and switch conditions (E2G, G2E). At stimulus onset, switch trajectories begin near onset point of prior task set and then evolves towards the repeat trajectories of the current task set. In **a,b**, each panel shows data from different subject-level models.

Gram–Schmidt orthonormalization ^(31)^ of the three semantic axes yielded an orthogonal matrix defining a subject-specific rigid transformation of each embedding (Methods). As the UMAP objective depends on the embedding only through pairwise distances, which rigid transformations preserve, UMAP embeddings are defined only up to such transformations; moreover, the alignment therefore reorients the embedded trajectories without altering any of their geometric relations. This transformation carried each subject-level UMAP embedding–whose native axes are determined by initialization and stochastic optimization and therefore carry no intrinsic identity or ordering ^(45)^–into a shared reference frame with labeled axes (Figs. 3a, 4; Extended Data Fig. 4b). Critically, this labeling is meaningful only to the extent that the raw semantic contrast vectors constitute an approximately orthogonal set within subject-level UMAP embeddings: orthonormalization leaves a near-orthogonal triad essentially unchanged, so each labeled axis remains closely aligned with the data-driven contrast vectors it is derived from, whereas strongly oblique contrast vectors would be displaced away from the directions they are meant to represent.

We therefore assessed the pairwise and joint orthogonality of the raw contrast vectors. The angular differences between the raw task type axis and either of the raw task-relevant target attribute axes were clustered around 90^◦^ across subjects: vs. gender axis, 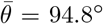 (circular V-test, *V* = 0.94, *r* = 0.95, *p*_Holm_ *<* 10^−3^); vs. emotion axis, 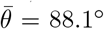 (*V* = 0.98, *r* = 0.98, *p*_Holm_ *<* 10^−3^). Although the two task-relevant target-attribute directions showed greater inter-subject variability, their angular difference was also aligned to 90^◦^ (Fig. 3b; 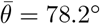, *V* = 0.54, *r* = 0.59, *p*_Holm_ *<* 10^−3^). Joint orthogonality was assessed by constructing a group-level Gram matrix across the three raw semantic contrast vectors (Methods), which showed diagonal entries near 1 and off-diagonal entries near 0 (Fig. 3d), consistent with an approximately orthogonal three-direction configuration. With respect to a null distribution constructed from random triads of unit vectors (Methods), the observed subject-averaged orthogonality error was smaller than expected by chance (Fig. 3e; observed = 0.61; null mean = 1.37, 95% interval (1.22, 1.52); *p <* 10^−3^).

Together, these results indicate that the semantic organization of latent states was not idiosyncratic to individual subjects, but instead reflected a shared representational configuration. Again these findings are specific to the UMAP embedding and need not carry over to the native 16D latent space as UMAP preserves local neighborhood relations rather than distances or angles (Methods).

Note that the emotion and gender attribute axes were defined from the corresponding repeat conditions in which those attributes were behaviorally relevant (E2E and G2G, respectively), indicating that those axes characterized latent dynamics with respect to not only the pure perceptual features of target stimuli, but also their behavioral relevance. To confirm this, we compared these axes with those obtained from repeat trials in which the same target attributes were task irrelevant. We found that the angular differences were not significantly aligned with 0^◦^ (Fig. 3c; gender: 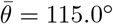, *V* = −0.23, *r* = 0.35, *p* = 0.94; emotion: 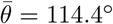, *V* = −0.25, *r* = 0.38, *p* = 0.95), indicating that the emotion and gender axes do not generalize across task types.

### 2.3 A hierarchical latent task representation

The UMAP embedding was used strictly as a visualization tool; aligning each subject’s embedding to a shared reference frame with labeled axes further enhanced the interpretability of the embedded latent dynamics, and motivated hypothesis-guided tests of their geometry. For example, visualization of repeat condition trajectories stratified by task type and all four combinations of gender and emotion attributes suggested a hierarchical organization (Fig. 5a). Again, we note that quantification and statistical testing of latent-state geometry in all subsequent analyses were performed in the original 16D latent space.

**Fig. 5:**
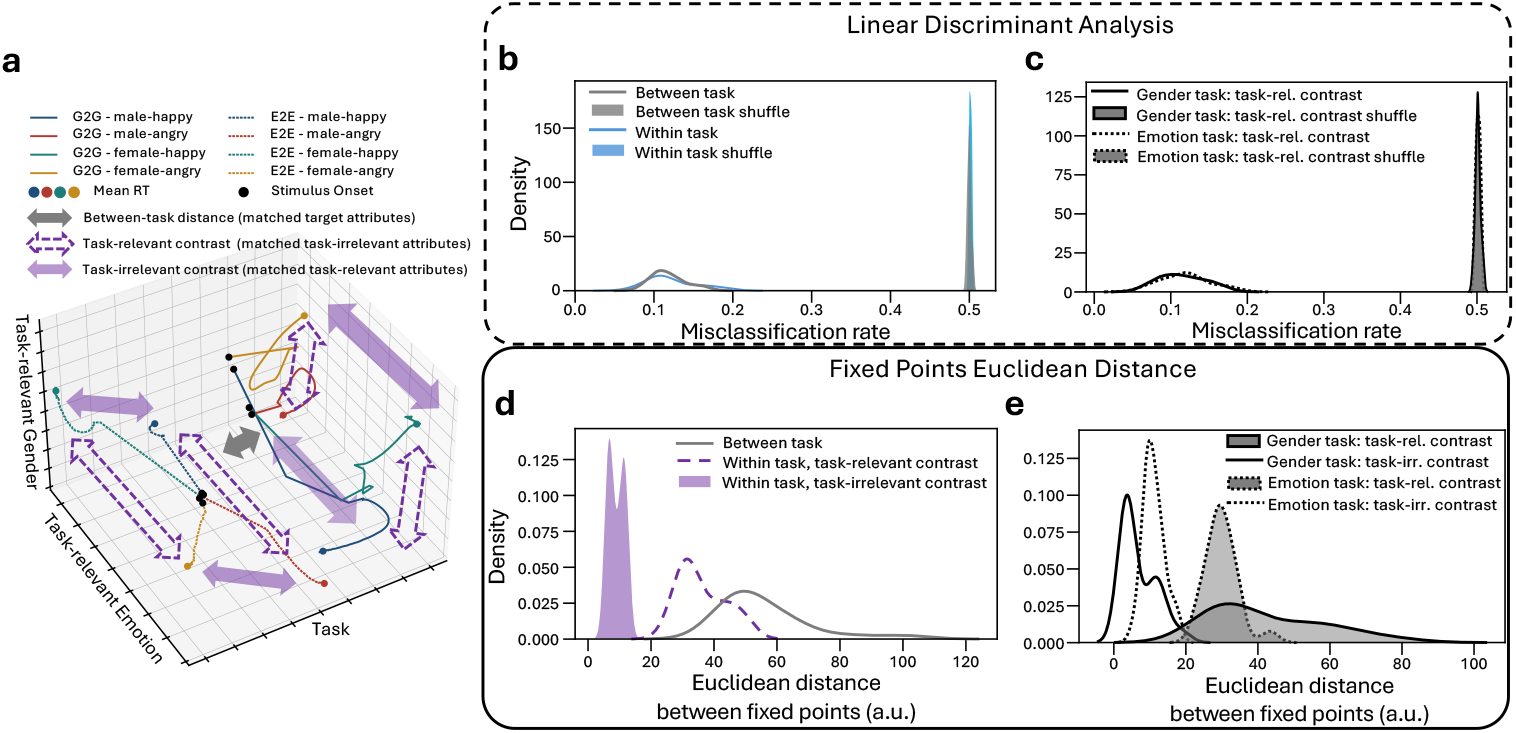
Hierarchical organization of repeat trajectories. **a**, Single subject example of trial-averaged repeat trajectories stratified across task type × emotion attribute × gender attribute. Arrows illustrate between-task type contrasts with matched target attributes (grey), task-relevant target attribute contrasts with matched task-irrelevant attributes (clear purple), and task-irrelevant target attribute contrasts with matched task-relevant attributes (solid purple). **b**, Separability between trajectories quantified using LDA misclassification rates at mean RT. **c**, LDA misclassification rates for task-relevant target attribute classification within each task controlled for the task-irrelevant attribute. **d**, Euclidean distances between fixed points. Between-task distances were computed across subsets of trials with matched gender and emotion attributes. Within-task distances were computed separately for task-relevant and task-irrelevant target attribute matching the other set of target attributes. **e**, Within-task distances shown separately for each task type. For **b**–**e**, *N* = 22 models.

To test for a hierarchical organization of repeat-condition trajectories, we quantified their separability using linear discriminant analysis (LDA). Taking the latent state along each trajectory at the time of the trial-averaged RT within the corresponding subset of trials, cross-validated LDA revealed robust separability at both the between- and within-task levels (Fig. 5b). Task type could be decoded with a misclassification rate of 0.12 ± 0.0045 (mean±s.e.m.), substantially lower than the corresponding label-shuffle null distribution (0.50 0 ±.0005; paired-sample *t*(21) = −85.75, *p*_Holm_ *<* 10^−26^). Within-task LDA was computed across all pairs of gender × emotion attribute contrasts, yielding a misclassification rate of 0.12 ± 0.0066, again substantially lower than the corresponding null distribution (0.50 ± 0.0003; paired *t*(21) = −57.60, *p*_Holm_ *<* 10^−23^). We found consistent results when task-relevant attribute classification was examined separately within the two task types while controlling for the task-irrelevant attribute (Fig. 5c). Misclassification rates were 0.11 ± 0.0062 for G2G and 0.12 ±0.0065 for E2E, compared with the corresponding label-shuffled null distributions (G2G: 0.50 ±0.0007, paired *t*(21) = −60.01, *p*_Holm_ *<* 10^−23^; E2E: 0.50 ±0.0006, paired *t*(21) = −58.78, *p*_Holm_ *<* 10^−23^).

The Euclidean distance between fixed points (Fig. 5d; Methods) similarly revealed a hierarchical organization. Between-task distances computed across subsets of trials matching gender and emotion attributes were largest (55.77±3.04 a.u., mean±s.e.m.), followed by within-task distances across task-relevant target attributes (matching task-irrelevant attribute; 36.16±1.57 a.u.), and finally by within-task distances across task-irrelevant target attributes (matching the task-relevant attribute; 9.01±0.51 a.u.). Distances across adjacent levels within this hierarchy were graded (task type vs. task-relevant attributes: paired *t*(21) = 11.00, *p*_Holm_ *<* 10^−9^; task-relevant vs. task-irrelevant attributes: paired *t*(21) = 23.29, *p*_Holm_ *<* 10^−15^). Consistent findings were obtained within each task type (Fig. 5e): in G2G, task-relevant gender contrasts produced larger fixed-point distances than task-irrelevant emotion contrasts (42.37 ± 3.30 vs. 6.93 ± 1.01 a.u.; paired *t*(21) = 15.04, *p*_Holm_ *<* 10^−11^); likewise, in E2E, task-relevant emotion contrasts produced larger distances than task-irrelevant gender contrasts (29.95 ± 0.93 vs. 11.09 ± 0.60 a.u.; paired *t*(21) = 35.49, *p*_Holm_ *<* 10^−19^).

Together, these findings demonstrate a hierarchical organization of repeat-trial representations, with the strongest separation across (i) task types, followed by (ii) task-relevant target attributes, and finally (iii) task-irrelevant target attributes.

### 2.4 Switch-repeat onset point distance tracks asymmetric switch costs

The hierarchical organization of repeat trajectories (Fig. 4a) was recapitulated by the early dynamics in switch trials, whereby trajectories began near the onset points of repeat trajectories from the previous task set (Fig. 4b, Extended Data Fig. 5a). The trajectories then traversed along the task-type axis toward the subspace of the current task set, ultimately converging with the repeat trajectories of the current task. We note that cumulative trajectory length from stimulus onset to each trial’s RT was strongly correlated with RT across trials within stimulus configurations (Extended Data Fig. 6), supporting the interpretation that traversal through latent space reflects the temporal evolution of within-trial behavior. Hence, we hypothesized that greater distance between the onset points of switch trajectories and the onset points of the repeat trajectories corresponding to the current task set would be associated with larger behavioral switch costs.

We defined onset point distance as the Euclidean distance–computed in the raw 16D latent space–between the onset points of switch trajectories (E2G and G2E) and corresponding repeat trajectories (G2G and E2E, respectively). Onset point distance was computed for each switch direction separately, averaging across subsets of trials with matched gender and emotion attributes (Fig. 6a; Extended Data Fig. 5b). We first confirmed that onset point distance was associated with behavioral switch costs across subjects (Figs. 6b,c): for E2G, Pearson’s *r* = 0.60 (bootstrap 95% CI (−0.16, 0.84), *p*_Holm_ = 0.0034); for G2E, Pearson’s *r* = 0.68 (bootstrap 95% CI (0.29, 0.86), *p*_Holm_ = 0.0011). We next asked whether this geometric metric recapitulated asymmetrically increased switch costs incurred when switching to the emotion task. Onset point distance was larger for G2E than for E2G (51.20 ±2.68 a.u. vs. 48.04 ± 2.45 a.u., mean s.e.m.; paired *t*(21) = −4.13, *p <* 10^−3^; Fig. 6d). Repeated-measures correlation ^(5)^ similarly showed that this association between onset point distance and behavioral switch cost across the two switch directions was consistent across subject-level models (Fig. 6e; *r*_rm_ = 0.66, bootstrap 95% CI (0.34, 0.84), *p <* 10^−3^).

**Fig. 6:**
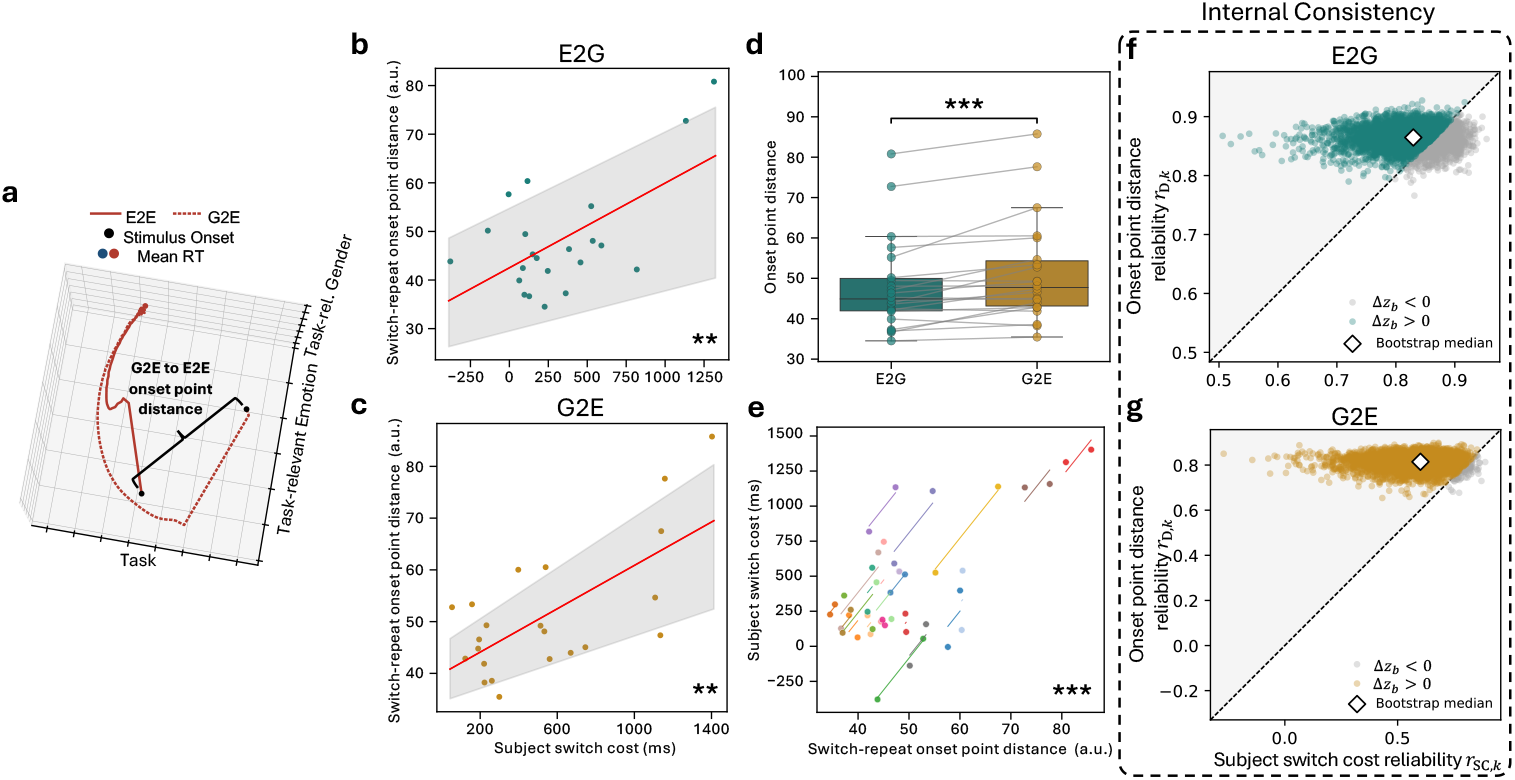
Switch-repeat onset point distance in latent space corresponds to behavioral switch costs. **a**, Illustration of the distance between onset points of switch trials and repeat trials of the current task type. The example shows the distance between the G2E and the E2E onset points. Metric computed as Euclidean distance in raw 16D latent space. **b,c**, Cross-subject correlation between raw switch costs and onset point distance for E2G (**b**) and G2E (**c**). Each circle represents individual subjects and the corresponding fitted model; red lines indicate linear fits; and gray shaded regions indicate 95% CI. **d**, Onset point distances for the two switch directions recapitulate the asymmetry in switch costs across directions. Each circle represents individual subjects, and gray lines connect paired E2G and G2E observations. **e**, Repeated-measures correlation between onset point distance and raw switch cost across the two switch conditions. Colors represent individual subjects, with lines connecting paired E2G and G2E observations depicted by circles. **f,g**, Internal consistency of onset point distance and raw switch costs were quantified across split halves for each switch direction separately. Reliability differences across these two metrics are shown for E2G (**f**) and G2E (**g**). Circles represent reliability results across paired trial-bootstrapped samples. Points along the dashed diagonal indicate equal reliability, whereas points above the diagonal indicate higher reliability of onset point distance (in 75.1% and 98.0% of bootstrapped samples for E2G and G2E, respectively). White diamonds indicate the bootstrap median reliability pair. ∗∗ *p <* .01, ∗∗∗ *p <* .001

Finally, we sought to assess the within-session reliability (internal consistency) of onset point distance in capturing individual differences in switch costs using a split-half design. To do so, trials were randomly partitioned into two non-overlapping halves; each half was modeled independently (Methods), and reliability was quantified with Spearman–Brown corrected cross-subject Pearson’s correlation across split halves for each switch direction separately (Figs. 6f,g). Onset point distance showed good reliability (*>* 0.75, ^(47)^) for both E2G (*r*_D_ = 0.873, 95% CI (0.825, 0.898), *p <* 10^−4^) and G2E (*r*_D_ = 0.822, 95% CI (0.743, 0.867), *p <* 10^−3^). Behavioral switch cost showed comparable reliability for E2G (*r*_SC_ = 0.878, 95% CI (0.706, 0.907), *p <* 10^−4^), but only moderate reliability for G2E (*r*_SC_ = 0.660, 95% CI (0.274, 0.788), *p* = 0.020).

For statistical comparisons of the reliability of the two metrics, we performed a paired trial-level bootstrapping within each subject × split halves while preserving the relative proportions of trials across task conditions (Methods). For each bootstrapped sample, each reliability measure was Fisher z-transformed, and their difference was computed as follows: Δ*z* = −*z*(*r*_D_) − *z*(*r*_SC_). The reliability of onset point distance was greater in 75.1% and 98.0% of bootstrapped samples in E2G and G2E, respectively. Statistical significance of these findings was assessed with respect to the bootstrapped distribution of Δ*z*: for G2E, onset point distance showed greater reliability than raw switch cost (Δ*z* = 0.369, 95% CI (0.022, 0.896), *p*_boot_ = 0.040; ^(46)^); for E2G, reliability was statistically indistinguishable (Δ*z* = −0.022, 95% CI (−0.229, 0.462), *p*_boot_ = 0.498).

Together, these results indicate that onset point distance provides a more reliable metric of individual differences in ATS performance, being favored over raw switch costs in the majority of bootstrapped samples for both switch directions, with a particularly strong reliability advantage for G2E.

### 2.5 Tortuosity of switch trajectory sub-trial segments corresponds to drift-diffusion model parameters

We next assessed correspondence between GeoDynFormer and a classical model of decision making, specifically the drift–diffusion model (DDM) ^(64;63)^. We fit RT distributions for each subject and task condition to compute the following DDM parameters: drift rate (*v*), which indexes the efficiency of evidence accumulation; non-decision time (*t*_0_), which captures processes outside evidence accumulation, including sensory perception, motor execution, and, in the case of switch trials, task-set reconfiguration. The fitted parameters demonstrated expected patterns across task conditions (Extended Data Fig. 7): non-decision time was longer for switch conditions for both task types, consistent with the behavioral cost incurred from task set reconfiguration ^(70)^.

We first confirmed that, across subjects, DDM parameters showed the expected correlations with trajectory length measured from the stimulus-onset point to the mean-RT point - a negative correlation between trajectory length and *v*, and a positive correlation with *t*_0_ in each condition (Extended Data Fig. 8). Building on these general relationships, we performed a more granular analysis of switch trajectories motivated by their visualizations in the common semantic embedding. As mentioned above, switch trajectories began in the subspace of the previous task set, which evolved to the subspace of the current task set first, prior to converging with the repeat trajectories of the current task set (Fig. 4b). In other words, the early segment of switch trajectories was consistent with non-decisional processes including task-set reconfiguration, whereas the late segment was consistent with evidence accumulation.

We therefore hypothesized that the geometry of trajectories during the early and late sub-trial epochs would show specific associations with DDM parameters capturing task-set reconfiguration (*t*_0_) and evidence accumulation (*v*), respectively. To do so, we partitioned each switch-trial trajectory using a transition point, *t*_*c*_, defined as the time point on the switch trajectory closest to the onset point of the repeat trajectory for the current task set (Fig. 7b; Methods): each trajectory was thereby segmented into early ([0, *t*_*c*_]) and late epochs ([*t*_*c*_, *RT*]). We then computed the tortuosity of each trajectory segment as 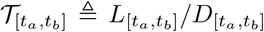 (over an interval [*t*_*a*_, *t*_*b*_]; ^(33;6)^), where 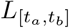 is the cumulative trajectory length and 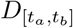 is the Euclidean distance between the start and end states of the corresponding epoch (Methods). Intuitively, tortuosity quantifies how indirect or geometrically inefficient a latent trajectory segment is and can therefore reflect the efficiency of task-set reconfiguration or evidence accumulation.

**Fig. 7:**
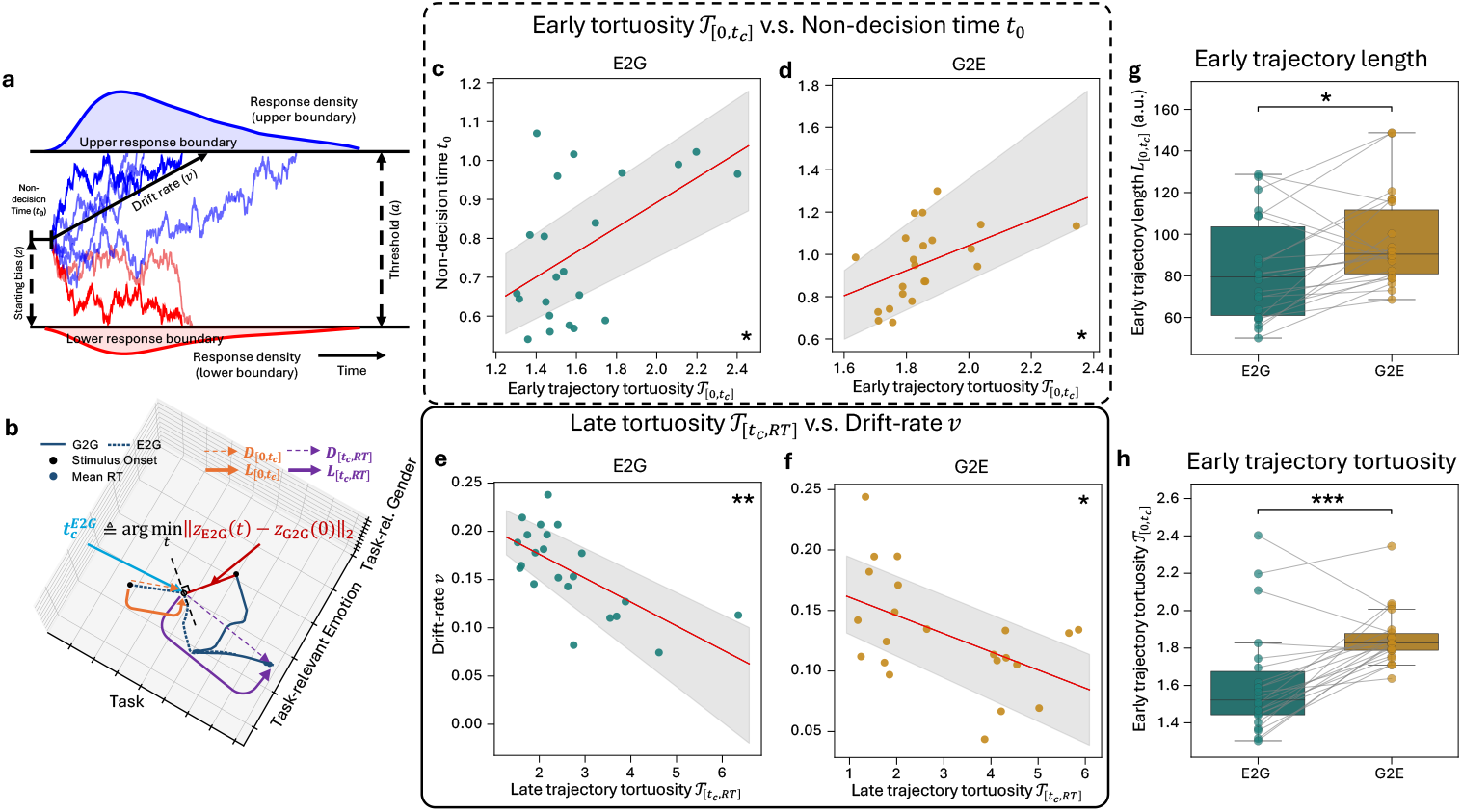
*t*_*c*_-based decomposition of switch-trial latent dynamics and DDM parameters. **a**, Schematic of the drift–diffusion model (DDM), illustrating drift rate (*v*), boundary separation (*a*), starting bias (*z*; fixed to *a/*2), and non-decision time (*t*_0_). **b**, *t*_*c*_-based decomposition of switch-trial trajectories, illustrated for E2G (dashed) and G2G (solid) trajectories. *t*_*c*_ is defined as the time point along the E2G trajectory closest to the G2G onset point, partitioning the E2G trajectory into early ([0, *t*_*c*_]) and late ([*t*_*c*_, *RT*]) epochs corresponding to sub-processes captured by DDM parameters *t*_0_ and *v*, respectively. For each epoch, trajectory length (*L*) is computed along the latent trajectory, whereas displacement (*D*) is the Euclidean distance between the start and end states. Trajectory tortuosity 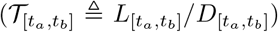 is then a geometric measure of the efficiency of each sub-trial epoch. **c,d**, *t*_0_ vs. early trajectory tortuosity 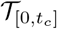 for E2G (**c**) and G2E (**d**) trials. **e,f**, *v* vs. late trajectory tortuosity 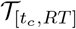 for E2G (**e**) and G2E (**f**) trials. **g,h**, Early trajectory length 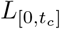 (**g**) and tortuosity 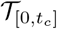 (**h**) for E2G and G2E switch trials. In **c**–**f**, circles represent individual subjects; red lines indicate best linear fits, and shaded regions indicate 95% CI. In **g,h**, circles represent individual subject, and gray lines connect paired values across switch directions. ∗ *p <* 0.05, ∗∗ *p <* 0.01, ∗∗∗ *p <* 0.001.

Indeed, we found that early trajectory tortuosity 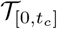 was positively correlated with non-decision time *t*_0_ (Fig. 7c, d; for E2G, Pearson’s *r* = 0.51, bootstrap 95% CI (0.15, 0.78), *p*_Holm_ = 0.03; for G2E, Pearson’s *r* = 0.49, bootstrap 95% CI (0.28, 0.72), *p*_Holm_ = 0.03). That is, subjects whose early switch trajectories were more tortuous tended to exhibit longer *t*_0_. By contrast, late trajectory tortuosity 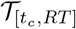 was negatively correlated with drift rate *v* (Fig. 7e, f; for E2G, Pearson’s *r* = −0.66, bootstrap 95% CI (−0.85, −0.49), *p*_Holm_ = 0.002; for G2E, Pearson’s *r* = −0.51, bootstrap 95% CI (−0.73, −0.24), *p*_Holm_ = 0.02)– i.e., subjects with more tortuous late-phase trajectories tended to exhibit lower *v*.

As early and late tortuosities depend on each other through *t*_*c*_, we sought to confirm that the above associations between sub-trial epoch tortuosities and DDM parameters held even when controlling for the tortuosity of the remaining epoch. For each switch direction, we fit multivariable regressions predicting *t*_0_ with 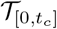 while controlling for 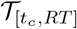, and likewise predicting *v* with _[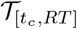_ while controlling for 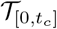 (Methods).

Nested model comparisons showed that the hypothesized geometric metric significantly improved fit: for *v*, 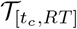 explained additional variance beyond early tortuosity in both E2G (Δ*R*^2^ = 0.330, *F* (1, 19) = 9.48, *p*_Holm_ = 0.0247) and G2E (Δ*R*^2^ = 0.218, *F* (1, 19) = 5.32, *p*_Holm_ = 0.0412); for *t*_0_, 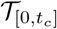 explained additional variance beyond late tortuosity in both E2G (Δ*R*^2^ = 0.264, *F* (1, 19) = 8.68, *p*_Holm_ = 0.0249) and G2E (Δ*R*^2^ = 0.173, *F* (1, 19) = 6.38, *p*_Holm_ = 0.0412). In the models for *t*_0_, 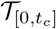 was confirmed to be the only statistically significant predictor in E2G trials (*β* = 0.51, *t*(19) = 2.95, *p*_Holm_ = 0.02; for 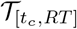, *t*(19) = 2.44, *p*_Holm_ = 0.07), while both 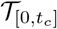 and 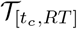 were predictors in G2E trials (*β >* 0.41, *t*(19) *>* 2.52, *p*_Holm_ *<* 0.05). In the models for *v*, 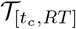 was the only statistically significant predictor in both switch directions (*β <* −0.46, *t*(19) *<* −2.30, *p*_Holm_ *<* 0.05; for 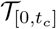, |*t*(19)| *<* 0.38, *p*_Holm_ = 1.0). Together, these results indicate that late trajectory tortuosity reflects drift rate, whereas early trajectory tortuosity reflects non-decision time.

Finally, we assessed whether geometric metrics captured the greater *t*_0_ observed in G2E relative to E2G trials (Extended Data Fig. 7). We found that early trajectory length 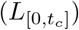 was greater for G2E trials (Fig. 7g; 96.71 ± 4.74 a.u. (mean ± s.e.m.) vs. 82.84 ±5.28 a.u.; paired *t*(21) = −2.77, *p* = 0.012). We found a similar difference in early trajectory tortuosity 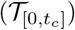 that was even more pronounced (Fig. 7h; G2E vs. E2G: 1.86 ±0.031 vs. 1.63 ±0.062, mean±s.e.m.; paired *t*(21) = −4.67, *p <* 10^−3^).

In sum, these findings show that an intuitive temporal decomposition of switch trajectories relative to *t*_*c*_ reveals the geometric correlates of sub-processes captured by DDM parameters. Early tortuosity tracked *t*_0_, whereas late tortuosity tracked *v*, indicating that the geometric inefficiency of switch-trial latent dynamics reflects the efficiency of the behavioral sub-processes underlying task-set reconfiguration and evidence accumulation.

### 2.6 Cross-condition angular alignment of trajectories reveals asymmetric flexibility of task sets

We next asked whether latent geometry could provide empirical support for one of the theories on the neurobehavioral origins of asymmetric switch costs in ATS. Schuch et al. ^(71)^ proposed that asymmetric switch costs arise because of differences in task set flexibility. The less flexible gender task is more shielded from the emotion task set during evidence accumulation, but has greater carryover effect (inertia) during task set reconfiguration in switching to the emotion task. In contrast, the more flexible emotion task is less shielded during evidence accumulation and has less inertia when switching to the gender task. While DDM parameter fits from our cohort were largely consistent with this account (*t*_0_ greater in G2E vs. E2G, and *v* greater in G2G vs. E2E; Extended Data Fig. 7), a more direct test would require comparing within switch and repeat trials across subsets of trials based on target attributes. In other words, the task set flexibility theory predicts that task-irrelevant target attributes should tax behavioral processes related to task set reconfiguration and evidence accumulation to a greater degree when performing the emotion task relative to when performing the gender task. However, DDM parameter fitting for such subsets of trials is precluded by insufficient trial counts, even for hierarchical implementations ^(76)^ (see Discussion).

To assess whether latent geometry could provide empirical support for the task set flexibility theory, we defined two geometric metrics based on angular similarity between latent trajectories (Figs. 8a,e; Extended Data Fig. 9): (i) a task-relevant alignment metric (*A*_rel_) quantified angular alignment between trajectories sharing the same task-relevant target attribute; (ii) the task-irrelevant alignment metric (*A*_irr_) quantified alignment between trajectories sharing the same task-irrelevant attribute. The instantaneous cosine similarity between trajectories was averaged across distinct subtrial epochs as follows: *A*_irr_ was computed during non-decision time ([0, *t*_0_]), whereas *A*_rel_ was computed during decision time ([*t*_0_, *RT*]). We hypothesized that the less flexible gender task would be associated with stronger alignment between trajectories with the same task-relevant target attribute when repeating tasks (i.e., *A*_rel_ greater for G2G vs. E2E)–consistent with stronger shielding–and that switching away from the less flexible gender task would incur stronger alignment between trajectories with the same task-irrelevant attribute (i.e., *A*_irr_ greater for G2E vs. E2G) - consistent with stronger inertia.

**Fig. 8:**
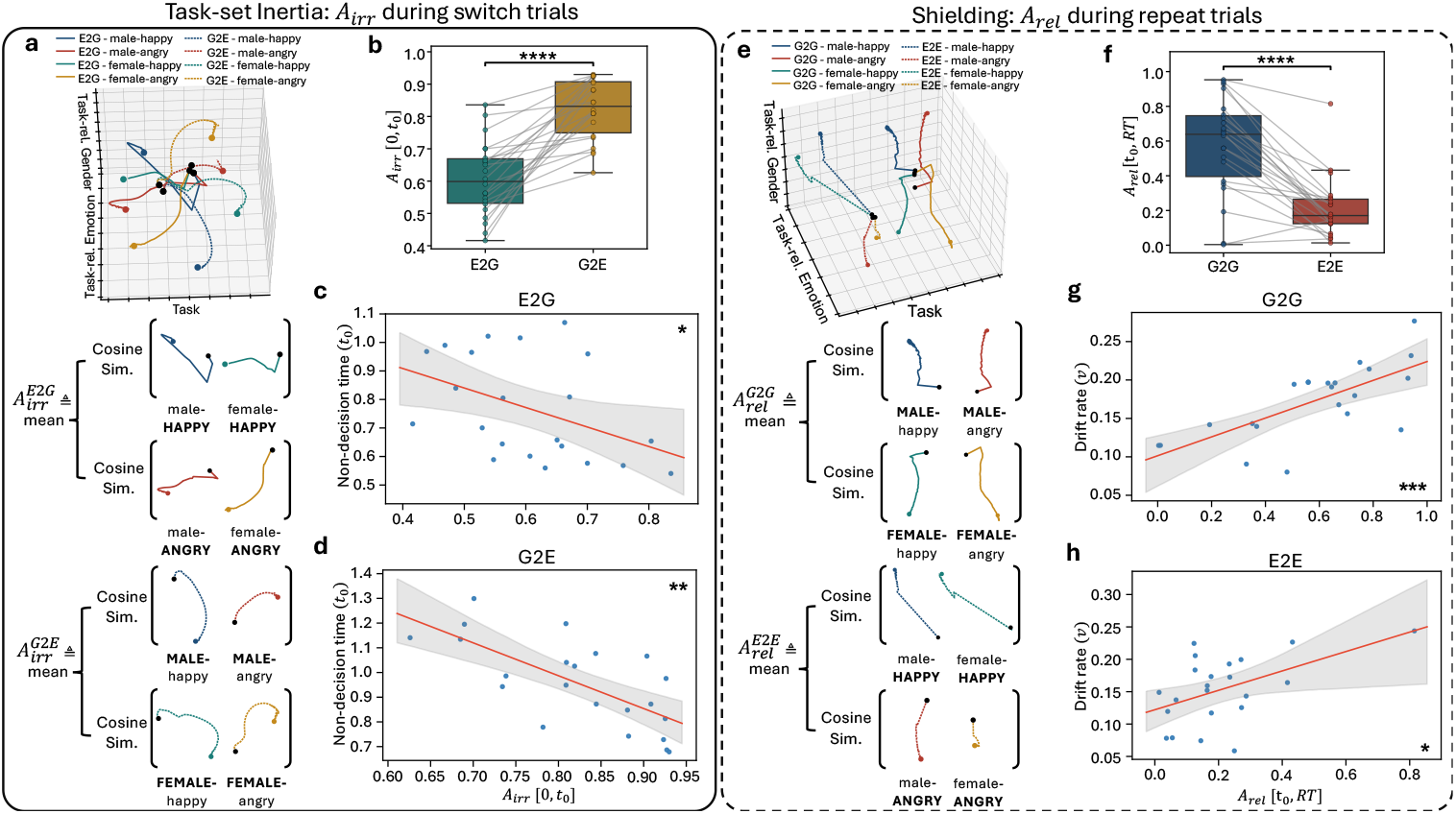
Trajectory alignment metrics reveal asymmetric flexibility of task sets. **a**, The task-irrelevant alignment metric *A*_irr_[0, *t*_0_] was computed from switch-condition trajectory pairs sharing the same task-irrelevant target attribute during non-decision time. **b**, *A*_irr_[0, *t*_0_] showed stronger alignment for G2E than E2G, indicating greater inertia of the gender task when switching away from it. **c,d**, Cross-subject correlation between *t*_0_ and *A*_irr_[0, *t*_0_] for E2G (**c**) and G2E (**d**) trials. **e**, The task-relevant alignment metric *A*_rel_[*t*_0_, *RT*] was computed from repeat-condition trajectory pairs sharing the same task-relevant target attribute during decision time. **f**, *A*_rel_[*t*_0_, *RT*] showed stronger alignment for G2G than E2E, indicating stronger shielding of the gender task set during repeat performance. **g,h**, Cross-subject correlation between *v* and *A*_rel_[*t*_0_, *RT*] for G2G (**g**) and E2E (**h**) trials. **a,e**, illustrate the alignment metrics with trajectories from example single-subject embeddings. In **c,d,g,h**, circles represent individual subjects, red lines indicate best linear fits, and shaded regions indicate 95% CI. ∗ *p <* .05, ∗∗∗ *p <* .001, ∗∗∗∗ *p <* .0001.

Indeed, we found that *A*_irr_ was greater for G2E (0.82±0.020) than for E2G (Fig. 8b; 0.61 ± 0.024; paired *t*(21) = −7.21; *p <* 10^−6^), indicating stronger inertia of the previous task set when switching away from the gender task. Moreover, *A*_rel_ was greater for G2G (mean±s.e.m.: 0.58 ±0.059) relative to E2E (Fig. 8f; 0.21 ±0.038; paired *t*(21) = 7.09; *p <* 10^−6^), indicating greater shielding from the emotion task set. We next asked whether these angular-alignment metrics would capture individual differences in DDM parameters. In switch trials, *A*_irr_ was negatively correlated with *t*_0_ across subjects (Fig. 8c,d; for E2G, Pearson’s *r* = −0.43, bootstrap 95% CI (−0.71, −0.07), *p*_Holm_ = 0.046; for G2E, Pearson’s *r* = −0.70, bootstrap 95% CI (−0.86, −0.45), *p*_Holm_ = 0.0010). In repeat trials, *A*_rel_ was positively correlated with *v* during both G2G (Fig. 8g; Pearson’s *r* = 0.70, bootstrap 95% CI (0.45, 0.89), *p*_Holm_ *<* 10^−3^) and E2E trials (Fig. 8h; Pearson’s *r* = 0.51, bootstrap 95% CI (0.05, 0.75), *p*_Holm_ = 0.011).

We further demonstrated the temporal specificity of these findings by testing the same associations outside of their hypothesized behavioral epochs. Switch-condition *A*_irr_ during decision time was not correlated with *t*_0_ (E2G: Pearson’s *r* = 0.04, bootstrap 95% CI (−0.37, 0.45), *p* = 0.86; G2E: Pearson’s *r* = 0.02, bootstrap 95% CI (− 0.39, 0.43), *p* = 0.94). Similarly, repeat-condition *A*_rel_ during non-decision time was not correlated with *v* (G2G: Pearson’s *r* = 0.26, bootstrap 95% CI (− 0.26, 0.53), *p* = 0.24; E2E: Pearson’s *r* = 0.08, bootstrap 95% CI (− 0.30, 0.51), *p* = 0.71).

Taken together, these findings provide support for the task set flexibility theory of asymmetric switch costs during ATS. Asymmetries in the angular alignment of repeat and switch trajectories based on task-relevant and irrelevant target attributes, respectively, were consistent with the gender task set being less flexible and the emotion task set being more flexible.

## 3 Discussion

Building on the backbone of dynamical VAE models independently fitted for each individual ^(39)^, GeoDynFormer casts task behavior as continuous stochastic trajectories through latent space, inferring sub-trial behavioral dynamics that are not directly accessible from the raw RT data. Relative to this backbone, we introduced an attention-based posterior network to better capture longer-range temporal context during inference, and adapted the fitting protocol so that subject-level models can be trained on datasets of typical laboratory scales. Interpretability was enhanced by visualizing trajectories in a low-dimensional embedding with labeled axes that were shared across independently trained subject-level models, which aided in characterizing the learned dynamics with a suite of intuitive geometric features. This geometric analysis pipeline, with metrics quantified in native latent space, resolved how ATS behavior is jointly shaped by task condition and individual differences. We demonstrate the potential of latent-geometry based metrics of task behavior to improve the reliability of task-based measurements of individual differences, to map theory-agnostic model dynamics onto theory-constrained model parameters, and to adjudicate theoretical accounts of task-based experimental effects (asymmetric switch costs in ATS).

### 3.1 Generative ANN models of individual subjects

Recent developments in ANNs and latent dynamical models have made it possible to investigate the multiscale temporal structure of task behavior with enhanced sub-trial temporal resolution. However, in the majority of ANN applications to study behavior, models are trained directly on the task, such that the resulting task-optimized latent dynamics of the model do not reproduce realistic behavior. Furthermore, models that are trained to fit behavior have commonly pooled data across multiple subjects ^(16;22;8;50)^, or have captured individual differences by coupling subject-specific components to group-trained ones. Examples include subject-specific student networks distilled from a teacher trained jointly on all subjects ^(40)^; subject-level embeddings inferred by a shared encoder-decoder trained jointly across all subjects ^(15;36)^; and per-subject fine-tuning initialized from a pre-trained group-level network ^(29)^. In sum, prior approaches leave individual differences either unparameterized or constrain subject-level estimates to group-derived teachers, manifolds, or initializations.

Most closely related to our approach, Jaffe et al. ^(39)^ fitted independent latent dynamical models (task-DyVA) to individual subjects during performance of a NTS paradigm. GeoDynFormer retains the dynamical VAE and state-space backbone while introducing the following modifications. First, future-conditioned posterior inference was implemented using a Transformer-based attention mechanism ^(75)^ in GeoDynFormer, rather than the backward long short-term memory (LSTM) network ^(37)^ used in task-DyVA, allowing temporally separated future observations to contribute directly to the inference context. In an otherwise matched architecture comparison across all subjects, Transformer-based inference yielded more accurate recovery of condition-wise RTs than LSTM-based inference (Extended Data Fig. 1). Second, whereas task-DyVA visualized latent trajectories using PCA, lower-dimensional visualization of latent trajectories in GeoDynFormer utilized a non-linear technique to prioritize local neighborhood relationships. We do not assume that UMAP itself is intrinsically more interpretable than PCA; rather, interpretability was further enhanced by extracting an approximately orthogonal set of semantic axes from each subject’s embedding, which allowed aligning each subject-level embedding to a shared reference frame with labeled axes. Third, our data augmentation method enabled training on substantially smaller subject-level datasets consisting of 240 trials relative to at least 5 hours of gameplay in task-DyVA (corresponding to approximately 10^4^ trials based on the reported inclusion criterion that trial-averaged RTs did not exceed 1.25 s). Fourth, model fitting in GeoDynFormer prioritized switch costs separately across the two switch directions, whereas task-DyVA used an aggregate switch cost along with a stimulus-feature congruency effect that was not applicable to our specific ATS paradigm. Whereas the first three modifications enhanced model performance and interpretability in ways that are expected to generalize to other behavioral paradigms, the fourth exemplifies modifications to the model training protocol specific to the behavioral paradigm of interest.

### 3.2 Task behavior measures based on geometry of latent dynamics

Geometric analysis of the GeoDynFormer model’s latent dynamics was facilitated by visualizing them as continuous trajectories in a lower-dimensional embedding ^(69)^. Whereas linear dimensionality-reduction methods preserve the geometry of the retained subspace exactly, a linear projection retains only the highest-variance directions and may therefore fail to visually separate structure residing in the remaining dimensions. We therefore used UMAP, which prioritizes preservation of local neighborhood relations (albeit at the expense of global metric fidelity) in the low-dimensional embedding ^(54)^.

Whether using linear or nonlinear methods, the intrinsic dimensions carry no intrinsic identity. We applied a rigid rotational transformation that enabled aligning subject-level embeddings to a common reference frame with labeled axes. To do so, we constructed an orthonormal basis within each subject-level embedding using contrast vectors between the fixed points of repeat-condition trajectories, stratified by task type, task-relevant emotion attributes, and task-relevant gender attributes–referred to as semantic axes. The three raw contrast vectors were subsequently Gram–Schmidt orthonormalized to define a subject-specific basis for alignment, allowing each subject’s UMAP coordinates to be re-expressed in a common reference frame. As the UMAP objective depends on the embedding only through pairwise distances, alignment to a common reference frame can be performed with any rigid rotation without altering the visualized geometric relations. Importantly, however, whether the axes within the common reference frame can retain the labels ascribed to the raw semantic axes depends on the degree to which they constitute an orthogonal set in each subject’s UMAP embedding. We demonstrated that the three raw contrast vectors between fixed points of emotion vs. gender repeat trials, angry vs. happy trials within E2E trials, and female vs. male trials within G2G repeat trials formed an approximately orthogonal set across subject-level UMAP embeddings. Our approach aligns with the broader use of dimensionality reduction and interpretable coding dimensions to study high-dimensional neural activity dynamics ^(52;7;12;60)^. Model interpretability is particularly important for applications in healthcare, where the utility of ANNs depends not only on predictive performance but also on whether their internal representations and outputs can be meaningfully understood by clinician end-users ^(41;38;49)^.

It is interesting to note that ATS-related variables captured by the semantic axes were either variables directly inputted into the model (task type) or their conjunctions (task type ×emotion attribute, task type ×gender attribute). In fact, whereas emotion and gender attributes were inputted into the model as pure sensory variables, the raw semantic axes constructed for task-irrelevant emotion and gender attributes were not aligned with corresponding task-relevant vectors.

A notable qualification is that UMAP is a nonlinear technique that does not preserve global distances or angles. Consistent with this limitation, the approximately orthogonal configuration observed among the raw semantic axes in subject-level UMAP embeddings was not present when the corresponding contrast vectors were recomputed directly in the original 16D latent space. Thus, the orthogonality of the three semantic axes should be interpreted as an organizational property of the UMAP embedding alone. The shared semantic embedding was therefore used primarily to enhance interpretation of the low-dimensional embeddings of latent trajectories and to motivate hypothesis-driven analyses of their underlying geometry.

For example, visualization of repeat-condition trajectories in the shared semantic reference frame suggested a three-tiered hierarchical organization of repeat-condition trajectories across task type, task-relevant target attributes, and task-irrelevant target attributes, which was statistically confirmed in the native 16D latent space. Switch trajectories began near the repeat-trial onset points of the previous task set, crossing over to the subspace of repeat trials of the current task set–motivating our analyses of sub-trial segmentations corresponding to the sub-processes captured by DDM parameters. Post-hoc analyses of the underlying geometry of latent trajectories were facilitated by visualizing their dynamics in the shared semantic embedding, leading to a suite of intuitive geometric metrics computed from single-condition trajectories (trajectory length, tortuosity) or from condition pairs (onset point distance and angular similarity) in the original 16D latent space. Future work should determine whether combinations of such geometric metrics can provide richer multivariate measures for behavioral phenotyping using task-based measurements.

### 3.3 Improving the reliability of task behavior

Task behavior is traditionally measured using trial-averaged summary statistics, such as condition-wise average RTs as well as their differences. In ATS, these measures have established robust group-level behavioral effects such as asymmetric switch costs, whereby switching to the affective task (e.g., face emotion judgments) incurs a greater cost vs. switching to the neutral task (e.g., face gender judgments) ^(66;21;20;48)^, as replicated in our study cohort. Whereas popular task designs are engineered to elicit robust group-level experimental effects, the same behavioral measures have been shown to be limited in their ability to capture individual differences across a wide range of behavioral paradigms ^(35;43;72;57)^. One of the contributory factors is their poor psychometric properties, specifically reliability, which can be quantified either within (internal consistency) or across sessions (test-retest reliability). Conceptually, reliability is decreased when error variance obscures the between-subject variance. There are several sources of error, including trial sampling noise, fluctuations in attention, state changes (e.g., mood), and trait changes.

Difference metrics generally have worse reliability vs. the two component metrics when they are of similar variance and are strongly correlated across subjects. Two mechanisms contribute. First, error variance propagates from both components into the composite; with independent errors, the error variances will accumulate ^(35;81;23)^. Second, subtraction removes any between-subject variance that is shared across conditions (e.g., overall RT). Such common signal cancellation can increase the proportion of error variance relative to the remaining between-subject variance ^(35)^.

We found that the distance between switch-condition onset points and the repeat-condition onset points of the current task set reflected both group-level effects shown with raw switch costs and correlated with individual switch costs. Critically, the onset point distance metric showed internal consistency comparable to or greater than raw switch cost across independently modeled, non-overlapping halves of the experimental session.

Notably, the reliability of raw switch costs has been reported to be relatively favorable in a similar implementation of ATS ^(20)^. Internal consistency was computed across 40 subjects using a split-half approach, with each half consisting of 48 repeat and 12 switch trials per task type: corrected Pearson’s correlation was 0.87 (95% CI 0.8-0.92) and 0.89 (0.83-0.94) for E2G and G2E, respectively. These results showing good reliability of raw switch costs suggest that subtracting repeat-trial RT from switch-trial RT isolates a distinct neurobehavioral process—task-set reconfiguration—specific to switch trials, mitigating the second mechanism mentioned above related to common signal cancellation, as predicted in Miller and Ulrich ^(57)^. However, raw switch costs remain susceptible to the first mechanism related to error propagation, as demonstrated by the sensitivity of raw switch cost internal consistency to trial numbers (i.e. measurement error) ^(20)^. In our study, internal consistency was computed across 22 subjects, with each split half consisting of approximately 10 repeat and 3 switch trials per task type. Consistent with the sensitivity of raw switch costs to trial sampling, we found that although internal consistency for E2G was comparable to numbers reported in ^(20)^ (*r*_SC_ = 0.878, 95% CI (0.706, 0.907)), internal consistency for G2E was substantially lower (*r*_SC_ = 0.660, 95% CI (0.274, 0.788)). In contrast, onset point distance showed good reliability (per benchmarks in ^(47)^) for both E2G (*r*_*D*_ = 0.873, 95% CI (0.825, 0.898)) and G2E (*r*_*D*_ = 0.822, 95% CI (0.743, 0.867)). Reliability of onset point distance was favorable vs. raw switch costs in the majority of bootstrapped samples for both switch directions, with a particularly strong and statistically significant advantage for G2E.

Although onset point distance introduces potential sources of variability not applicable to raw switch costs (e.g., stemming from differences in the models trained independently for each split-half), these results indicate that any influence from additional sources was outweighed by the reduced sensitivity of onset point distance to trial sampling noise. One possible explanation for this is that onset point distance is determined by latent dynamics estimated across the broader half-session, whereas raw switch costs are a direct function of trial-averaged RTs for each condition, and may therefore be more sensitive to reduced trial counts particularly for the switch condition. Along these lines, the disproportionate impact of trial counts on the reliability of raw switch costs for the G2E condition, compared with the relatively similar reliabilities of onset point distance across the two switch directions, may reflect the heavier tail observed in G2E RT distributions ^(20)^. Future work should assess the reliability of other geometric metrics introduced in this study, identify differential sources of measurement variability across geometric metrics, and determine whether multivariate combinations of geometric features lead to further gains in reliability.

### 3.4 Probing theoretical accounts of asymmetric switch costs in ATS

Classical accounts of asymmetric switch costs developed from NTS paradigms are anchored on the notion of task set “dominance.” A classic NTS paradigm is the Stroop color/word task, in which subjects sample color words (e.g., “red”, “blue”) printed in ink color that is either congruent (e.g., “red” printed in red) or incongruent with the word itself (e.g., “red” printed in blue). Switching to the word-reading task incurs greater switch costs vs. switching to the color-naming task ^(3;2)^. Here, the word-reading task is considered to be the more dominant task set due to the inherent automaticity of word-reading vs. color naming, leading to differential behavioral strengths in their performance. Accordingly, dominance has been operationalized based on two criteria. First, the dominant task set is associated with faster performance when performed in isolation (single-task/pure blocks) ^(3;2;79)^ or when repeating ^(77;79)^. Consistent findings have been reported when dominance of a task set is manufactured with recent practice ^(78)^ or is related to performing the task in their preferred language in bilingual individuals ^(56)^. Second, the dominant task set exerts greater cross-task interference while being less susceptible to it, such that behavioral slowing when responding to incongruent bivalent stimuli relative to univalent stimuli is smaller for the more dominant task ^(3;79)^. Here, for word reading, examples of incongruent bivalent and univalent stimuli would be “red” printed in blue and “red” printed in black, respectively; for color naming, “red” printed in blue and “xxxxx” (a string not related to colors) printed in blue, respectively. Two theories have been proposed linking task set dominance to asymmetric switch costs. First, switching to the more dominant task requires overcoming stronger backward inhibition to re-activate the task (negative priming): this was evidenced by findings that after color-naming, word-reading was slower for not only bivalent ^(77)^ but also univalent stimuli ^(2)^. Second, it has been proposed that the higher degree of cognitive control expended to suppress the more dominant task set may also carry over when repeating the less dominant task (endogenous control theory), leading to its faster performance (positive priming) ^(79)^.

In ATS, however, neither criterion applies to the task set associated with higher switch costs (emotion task). First, average RT has not been shown to be any different between the two tasks when performed in isolation ^(66)^ or when repeating ^(20)^. Similarly, in our cohort, there was no RT difference across tasks when performed in isolation, and the emotion task actually had greater RTs vs. the gender task when repeating. Hence, by the criteria developed based on NTS paradigms, the emotion task would be considered the less dominant task, and yet switching to the emotion task incurs greater switch costs. Second, as stimuli used in ATS paradigms (emotional faces) are inherently bivalent, a true univalent control is poorly defined. Cross-task interference has been operationalized in alternative ways for ATS paradigms, although findings point towards the opposite direction, i.e., that the non-affective task is the more dominant task set. First, in single-task blocks, variable face gender prolonged emotion judgments whereas variable face emotion did not affect the speed of gender judgments, indicating greater interference exerted by the gender task set ^(4)^. Second, switching to non-affective tasks (age or gender judgments) were decreased when mixed with an affective task (emotion judgment), but not when mixed with the other non-affective task, indicating less interference exerted by the emotion task ^(71)^.

In sum, although switching to the emotion task incurs a greater behavioral cost, the abovementioned theories based on differential negative and positive priming are anchored on the notion of task set dominance that does not suggest the emotion task to be the more dominant task set with the criteria used in NTS paradigms. Hence, it has been proposed that the emotion task may be considered to be more dominant based on distinct neurobehavioral processes related to the inherent bottom-up salience of emotional information/features. Consistent with this, neuroimaging studies implicate overlapping but distinct networks of brain regions whose activation correlates with task switching behavior across neutral and affective task versions ^(66;21;44;80)^. Moreover, individual differences in the underlying construct–AF–have been consistently related to self-reported measures of psychological resilience ^(61;28;51)^, a protective factor across the psychiatric spectrum. Converging evidence comes from studies that show the relationship with resilience is greater for AF vs. affectively-neutral cognitive flexibility ^(55;61;28)^, as well as vs. affective and neutral versions of the remaining components of executive function ^(58)^–updating working memory and inhibition ^(55)^.

Schuch et al. ^(71)^ offered an alternative account anchored on the notion of task set flexibility, proposing that the emotion task constitutes the more flexible task set, with less inertia across trials and less shielding within trials. In contrast, the gender task is the more inflexible task set, with greater inertia and stronger shielding. Our geometric analysis provided empirical support for this theory: angular trajectory alignment was greater between gender repeat trials sharing the same task-relevant gender attributes vs. emotion repeat trials sharing the same task-relevant emotion attributes, consistent with stronger within-trial shielding of the gender task set; alignment across G2E switch trials sharing the same task-irrelevant gender attribute was greater vs. E2G switch trials sharing the same task-irrelevant emotion attribute, consistent with greater cross-trial persistence of the gender task set.

### 3.5 Synergy with sub-process models

Drift–diffusion models (DDMs) ^(64;63)^ decompose task behavior into sub-trial processes involved in evidence accumulation and non-decisional components (stimulus perception, motor execution, and task set reconfiguration).

Our results link theory-agnostic descriptive/statistical models with more mechanistically-constrained models in two ways. First, motivated by the non-specific finding that condition-wise trajectory lengths positively correlated with non-decision time and negatively correlated with drift rates, we identified more specific geometric substrates of DDM parameters during switch trials. To do so, we defined sub-trial epochs based purely on the model’s internal dynamics, with respect to *t*_*c*_, defined as the point along the switch trial trajectory that was closest to the repeat trial onset point of the current task set. We showed a specific relation between the tortuosity of switch trajectories during the early vs. late epochs and DDM parameters indexing non-decision processes and evidence accumulation, respectively.

Second, DDM parameters were utilized in conjunction with geometric metrics to provide support for the task-set flexibility theory of asymmetric switch costs. Angular similarity of repeat trajectories stratified across task-relevant target attributes was greater in the G2G condition, but only when computed during later epochs corresponding to decision time (after *t*_0_); angular similarity of switch trajectories stratified across task-irrelevant target attributes was greater in the G2E condition, but only when computed during early segments corresponding to non-decision time (before *t*_0_).

Note that this analysis disentangled the behavioral effects of task type × task transition, but also further stratified by target attribute combinations (angry/happy × male/female). Estimating these differences with DDM parameters is precluded by the insufficient number of repeated trials per condition. Even with a hierarchical DDM implementation ^(76)^, the detection rate for differences in DDM parameters (based on simulated data with a twofold difference) was only 0.6 when computed with 75 trials per condition and 20 subjects (see Fig.8 in ^(76)^). In contrast, statistical differences in angular similarity were shown with 50 trials per condition (comparing repeat trials stratified across relevant target attributes) or 12 trials per condition (comparing switch trials stratified across irrelevant target attributes), highlighting the superior statistical power of latent geometry-based metrics of task behavior.

## 4 Methods

### 4.1 Description of Affective Task-switching Paradigm

Affective flexibility was assessed using an established affective task switching (ATS) paradigm in which subjects switch between an affective and affectively-neutral task while sampling naturalistic face images. Each task involved making binary judgments of the face’s emotion (happy vs. angry; emotion task) or the face’s gender (male vs. female; gender task) ^(21;20;48)^.

In our implementation (Figs. 1a,b), we used unique-identity faces from the FACES ^(19)^ database. Subjects first performed 2 single-task blocks comprised of 20 trials of either the gender or emotion tasks (pure trials). Subjects performed 4 mixed blocks each comprised of sequences of 3-7 repeats of the same task followed by a switch trial. This yielded four trial types: gender repeat (G2G), emotion repeat (E2E), gender switch (E2G), and emotion switch (G2E). Trials in each condition were balanced across target emotion and gender attributes. Subjects completed 240 mixed block trials: 96 repeat trials per task; 24 switch trials per direction. Each trial began with a fixation cross (1 s), followed by a scrambled-face (1.5–2 s), and then simultaneous presentation of the face target along with a colored border that indicated the current task set (cue) (Fig. 1a). Hence, in our case, the cue-target interval was 0, which maximizes switch cost asymmetry ^(70)^. Subject response (or if subjects failed to respond within 2 s) was followed by a blank screen (1 s). Experiments utilized Psychtoolbox ^(10)^

to present visual stimuli and collect behavioral responses.

Behavioral data were obtained from 22 subjects (*N* = 7 F, *N* = 15 M; mean age ± SD, 38.8 ±9.6 years; range, 22 −56 years). Subjects performed the task well, with overall accuracy 94.82% ±1.02% (repeat trial accuracy 95.41% ±1.02%; switch trial accuracy 92.50% ±1.62%). To verify above-chance performance at the individual-subject level, one-sided one-sample *t*-tests were performed on trial-wise accuracy against the 50% chance level, separately for repeat and switch trials. Performance was significantly above chance for all 22 subjects for both repeat and switch trials (subject-level *p <* .025).

### 4.2 The Affective Task Representation in GeoDynFormer

#### 4.2.1 Stimuli and response representation

To train GeoDynFormer, we converted the behavioral data from the affective task into a compact time-varying representation. Time was discretized into 20-ms steps, and at each step a set of binary-valued units indicated the presence or absence of the two task cues, two gender attribute values, and two emotion attribute values (Fig. 1c, left). This representation provided the model with the moment-by-moment task-cue and face-target information available on each trial, which in turn drove the evolution of the latent state through the transition dynamics (Eq. (1)).

Each trial was input from stimulus onset until the observed behavioral response time (RT) extended by 500 ms to accommodate cases in which the model-generated response occurred later than the subject’s actual response. To introduce variability during training, independent zero-mean Gaussian noise (s.d. = 0.1) was added to each input unit, with the noise resampled at every training iteration.

The observed response was represented as a smooth time-varying signal centered on the subject’s RT. Specifically, for each trial we placed a Gaussian kernel (s.d. = 300 ms, peak = 1) at the observed RT, producing a continuous response template for model fitting (Fig. 1c, right). This formulation allowed the model to learn both response identity and response timing within a unified temporal framework.

#### 4.2.2 RT calculation

After training, the model typically produced a temporally localized activation profile in one of the four response-output channels on each trial (Fig. 1c, right). To determine the model’s chosen response, we identified the output unit with the strongest activation within a time window beginning 100 ms after stimulus onset and ending at stimulus offset.

The model RT for a given trial was defined as the earliest time point at which the cumulative activation of the selected output unit reached one-half of its total activation within that window:

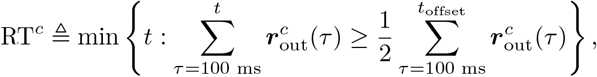

where 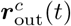 denotes the activation of the selected output unit at time *t*, and *t*_offset_ denotes stimulus offset. In other words, the model RT was defined as the temporal centroid of the chosen response profile.

Trials were excluded from RT estimation when the effective response window was truncated before a stable response profile could be established.

### 4.3 The GeoDynFormer Framework

#### 4.3.1 Generative model

The generative component of GeoDynFormer, which is responsible for emulating subject behavior, is a dynamical latent-variable model that takes sequences of task stimuli, ***s***_1:*T*_, as inputs and generates sequences of task responses, ***r***_1:*T*_, as outputs. The formulation is broadly related to dynamical variational autoencoder frameworks ^(30)^ and follows the general state-space formulation used in task-DyVA ^(39)^.

We define the latent state variables ***z***_1:*T*_ to evolve according to locally linear transition dynamics:

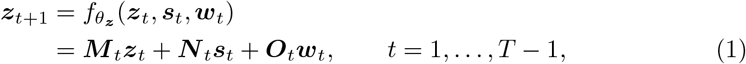

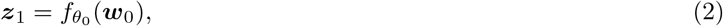

where ***w***_*t*_ denotes a stochastic latent perturbation and 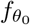 is an initialization function parameterized by a two-layer multilayer perceptron (MLP; hidden layer: 64 ReLU units ^(1)^; output layer: 16 linear units).

After training, the stochastic variables ***w***_*t*_ are sampled from a learned Gaussian prior,

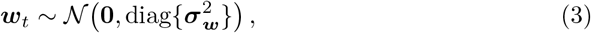

where the prior variance vector 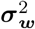 is learned. During training, by contrast, these variables are sampled from the encoder model.

To preserve interpretability while retaining flexibility, the transition matrices ***M*** _*t*_, ***N*** _*t*_, and ***O***_*t*_ are expressed as time-varying linear combinations of a small set of learned time-independent matrices:

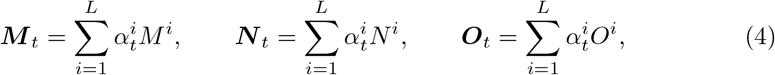

where the same weight vector ***α***_*t*_ is used for all three matrix families and is computed by a linear network 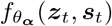 with softmax output units. In our implementation, ***z***_*t*_, ***w***_*t*_ ∈ ℝ^16^, ***s***_*t*_ ∈ ℝ^6^, and ***r***_*t*_ ∈ ℝ^4^, with *L* = 2 denoting the number of learned time-independent basis matrices used to construct each time-varying transition matrix.

The model outputs ***r***_*t*_, which correspond one-to-one with the possible task responses, are generated by the decoder model according to

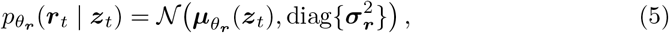

where 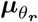 is parameterized by a two-layer MLP (hidden layer: 64 ReLU units; output layer: 4 sigmoid units). For response generation on the validation and test sets, the decoder output at each time point was taken as the conditional mean, 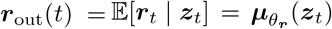 rather than sampling from 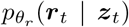. The decoder standard deviation vector ***σ***_***r***_, which was required only for evaluating the training objective, was fixed to 0.75 in all dimensions.

#### 4.3.2 Encoder model

To learn the generative model parameters, we used a variational autoencoder (VAE) framework ^(67)^ to perform approximate inference over the stochastic variables ***w***_0:*T* −1_. Because the conditional likelihood *p*_*θ*_(***r***_1:*T*_ | ***s***_1:*T*_) is intractable owing to the nonlinear transition dynamics (Eq. (1)) and decoding functions, we introduced an encoder model, *q*_*ϕ*_(***w***_0:*T* −1_ | ***s***_1:*T*_, ***r***_1:*T*_), which serves as an approximate posterior over the latent stochastic variables given the observed stimulus and response sequences. Following prior work on dynamical VAEs ^(42)^, we defined the approximate posterior over ***w***_0:*T* −1_, rather than directly over ***z***_1:*T*_, so that the latent state trajectory evolves deterministically once ***w***_0:*T* −1_ is specified. The encoder is used only during training and does not contribute to response generation after learning is complete.

The approximate posterior was factorized as

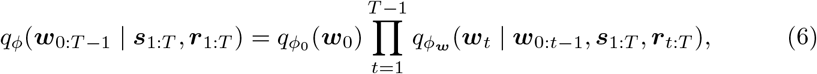

which preserves the overall sequential structure of the exact posterior ^(26;30)^ while introducing additional dependencies for computational convenience. In particular, the encoder conditions on the full stimulus stream and future response observations, allowing information from later time points to inform inference at the current time step.

Future observations were encoded into a context representation using Transformer-based attention ^(75)^. The resulting context variable *h*_*t*_ was combined with the current latent state *z*_*t*_ to parameterize the posterior over *w*_*t*_:

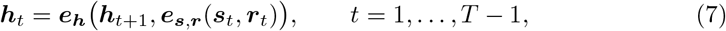

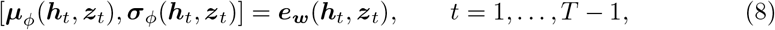

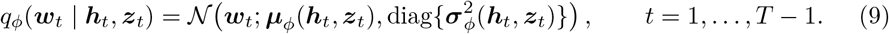

The latent state ***z***_*t*+1_ is then updated using the same transition dynamics as in the generative model, except that ***w***_*t*_ is sampled from the encoder rather than from the prior.

In our implementation, ***e***_***s*,*r***_ was parameterized by a two-layer MLP (hidden layer: 64 ReLU units; output layer: 64 linear units). The context encoder ***e***_***h***_ was implemented as a single Transformer layer with 64 hidden units and 4 attention heads, applied to the positionally encoded concatenated [***s***_*t*_, ***r***_*t*_] representation. Both ***µ***_*ϕ*_ and ***σ***_*ϕ*_ were implemented using single-hidden-layer MLPs with a shared 64-unit ReLU hidden layer; the output layer of ***µ***_*ϕ*_ contained 16 linear units, whereas that of ***σ***_*ϕ*_ contained 16 softmax units. The resulting variance values were multiplied by a fixed factor of 16 and offset by 10^−6^ for numerical stability. The model was initialized by sampling ***w***_0_ from the prior in Eq. (3), so that 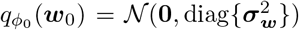, and ***z***_1_ was then obtained deterministically from 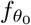.

#### 4.3.3 Training Objective

Model parameters were learned by maximizing a modified evidence lower bound (ELBO), the standard variational objective used to train variational autoen-coders by optimizing a lower bound on the conditional likelihood. To obtain this objective, we first derived the conditional density *p*_*θ*_(***r***_1:*T*_, ***w***_0:*T* −1_ | ***s***_1:*T*_) by integrating out the deterministic latent state variables ***z***_1:*T*_ from the full joint distribution of the generative model, following ^(30)^.

Conditioned on the stimulus sequence, the complete joint density over observed responses, stochastic variables, and latent states can be written as

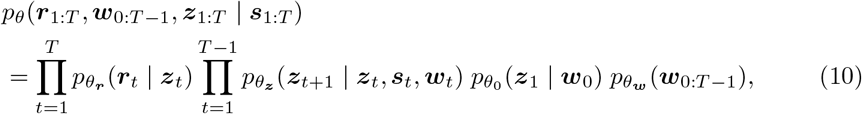

where the factorization follows directly from the state-space assumptions of the generative model. Because the latent transition is deterministic given ***z***_*t*_, ***s***_*t*_, and ***w***_*t*_, the transition densities can be rewritten using Dirac delta distributions:

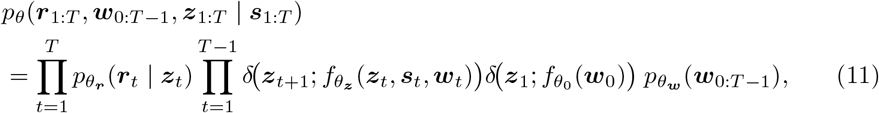

To eliminate ***z***_1:*T*_, we define a deterministic sequence **d**_1:*T*_ recursively by

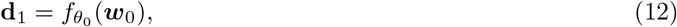

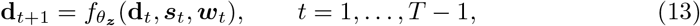

so that ***z***_*t*_ = **d**_*t*_ under the deterministic transition dynamics. Substituting **d**_*t*_ into Eq. (11) and integrating out the latent states sequentially yields

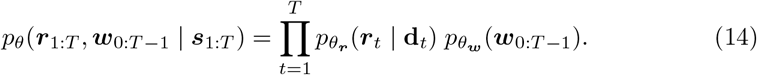

The resulting training objective was an annealed ELBO:

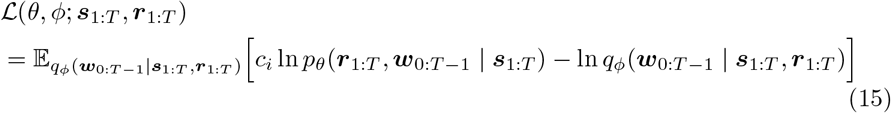

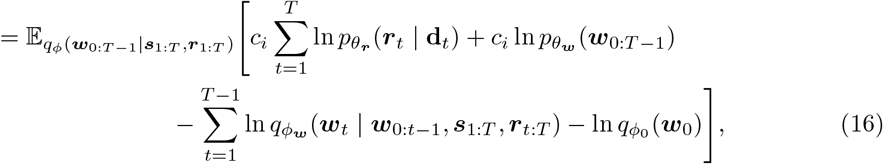

where Eqs. (6) and (14) were substituted into Eq. (15).

This objective differs from the standard ELBO in that the joint log-likelihood term is scaled by an annealing coefficient, 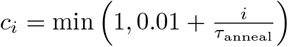, which increases monotonically with the number of gradient updates *i*. This annealing schedule improved training stability by preventing the reconstruction term from dominating too early in optimization, thereby allowing the posterior and generative dynamics to co-adapt more gradually. For all models, we set *τ*_anneal_ = 40000.

#### 4.3.4 Transformer-based posterior inference

The principal architectural distinction of GeoDynFormer lies in its posterior-inference model. As in Task-DyVA ^(39)^, inference is future conditioned: observations from the current and subsequent time points are used to construct a context representation that, together with the forward-evolving latent state, parameterizes the posterior over the stochastic transition variables. Task-DyVA implements this context encoder using a backward LSTM, such that information from future observations is propagated recursively through the recurrent hidden state. In contrast, GeoDynFormer uses Transformer-based attention ^(75)^, allowing temporally separated observations to contribute directly to the future-conditioned context representation. This distinction is particularly relevant for behavioral paradigms whereby trial history impacts behavior on the current trial. For instance, in task switching paradigms, switch vs. repeat status is not explicitly supplied to the model and must instead be inferred from the temporal history of task cues, stimuli, and responses.

To directly evaluate whether the Transformer-based inference architecture provided an empirical advantage, we trained a matched task-DyVA comparison model for each subject in which the Transformer context encoder was replaced by the recurrent LSTM-based inference architecture of Jaffe et al. ^(39)^. The comparison models were trained on the same subject-specific ATS datasets using the same stimulus and response representations, train–validation–test splits, data augmentation, generative-model configuration, training objective, optimization settings, and behavioral early-stopping and checkpoint-selection procedure as GeoDynFormer. Thus, the comparison was designed to isolate the posterior-inference architecture while holding the remainder of the ATS modeling framework fixed. Transformer-based inference provided more accurate recovery of condition-dependent reaction times than recurrent inference (Extended Data Fig. 1a).

#### 4.3.5 Configuration for latent-state interpretation

To visualize the temporal evolution of condition-dependent latent states, we projected the learned latent dynamics into a three-dimensional UMAP embedding ^(54)^. Whereas linear dimensionality-reduction methods capture global linear structure, UMAP was used here to preserve local neighborhood structure and provide a low-dimensional view of potentially nonlinear organization in the learned latent dynamics.

More broadly, this configuration was designed to facilitate model understandability by making internal state evolution directly examinable and relating its presentation to domain-relevant task variables, consistent with broader principles of interpretable and understandable machine learning ^(68;41)^. Here, behavior evolves through an explicit time-varying latent state, allowing the model’s within-trial dynamics to be examined directly as trajectories through state space. Because the native UMAP coordinates themselves have no intrinsic task-related meaning, we further constructed subject-specific semantic axes from condition-specific fixed points and used these axes to re-express each subject’s embedding in a common task–task-relevant emotion attribute–task-relevant gender attribute reference frame (Sections 4.5.1 and 4.5.4). Together, the latent dynamical representation, trajectory visualization, and semantic alignment provided an interpretable scaffold for examining how internal task representations evolved across conditions and for generating geometric hypotheses about the underlying dynamics.

Importantly, the UMAP embedding and semantic reference frame were used primarily for visualization and hypothesis generation rather than as substitutes for the learned latent representation itself. Except where explicitly noted, all quantification and statistical testing of geometric metrics were performed directly in the original 16-dimensional latent space.

### 4.4 Model Training

#### 4.4.1 Training, validation, and test splits

Each subject’s behavioral data were randomly assigned to three non-overlapping splits: 70% for training, 10% for validation, and 20% for testing. The validation split was used to monitor model performance during optimization and to select the retained model checkpoint, and was not used for parameter updates. The matched posterior-inference architecture comparison shown in Extended Data Fig. 1 was also evaluated on the validation split, using correct trials only. All other model-performance and latent-state analyses reported in the main and Extended Data figures were performed using correct trials from the held-out test split, which remained separate from both model training and checkpoint selection. Analysis-specific correctness criteria for transition-dependent measures are described in the corresponding subsections.

#### 4.4.2 Data augmentation

As our goal was to develop a modeling framework compatible with typical laboratory-scale behavioral datasets, we increased the effective number of training sequences through overlapping-window preprocessing and data augmentation. After extracting and filtering trial-level behavioral data (see Section 4.1), each subject’s experimental session was first segmented using a sliding window of 50 trials with stride 1. This procedure generated overlapping trial subsets while preserving the local temporal ordering of task stimuli, responses, and RTs, which served as the basis for constructing the temporally segmented training sequences described below.

We next applied data augmentation to increase the amount of training data and reduce imbalances between repeat and switch trials. The augmented training corpus consisted of a mixture of observed trial sequences and synthetic sequences generated by resampling RTs from the subject’s observed condition-specific distributions. For this procedure, behavioral data were segmented into 5-s sequences, corresponding to approximately 2-4 trials, and RT probability density functions were estimated separately for each of the four conditions (G2G, E2E, E2G, and G2E) using Gaussian kernel density estimation (KDE; bandwidth parameter = 0.25). Sequences containing extreme RT outliers were excluded before estimating these distributions (see Section 4.4.3).

To expand the training corpus, the set of 5-s training sequences was replicated tenfold. For each resulting sequence, we stochastically determined whether it was retained unchanged or replaced by a synthetic sequence. Observed sequences were retained with probability 0.25, whereas synthetic sequences were generated with probability 0.75. Observed sequences were retained with their originally recorded responses, including behavioral errors, whereas synthetic sequences were generated with the correct task response.

To generate a synthetic trial, one of the four conditions (G2G, E2E, E2G, or G2E) was sampled with equal probability to the observed set of trials, and an RT was randomly drawn from the corresponding condition-level KDEs. Face gender and emotion were then sampled independently with equal probability across the four gender×emotion target-attribute combinations. This resampling procedure was repeated until the cumulative duration of the sequence exceeded 5 s. The resulting sequence was then converted into the continuous time-varying representation inputted into GeoDynFormer.

Synthetic trials were assigned the correct task response rather than reproducing the subject’s trial-wise error distribution. Accordingly, the modeling objective focused primarily on reproducing subject-specific response timing and its variation across task conditions rather than individual differences in response accuracy. Because synthetic sequences constituted the majority of the augmented training corpus and contained only correct responses, fitted-model accuracy was at ceiling. Hence, the models were not intended to reproduce subject-specific error behavior.

The 5-s training duration was chosen to retain several consecutive trials within each sequence, thereby preserving the local cross-trial context needed to infer repeat and switch structure while maintaining a large number of training sequences and tractable optimization. Because sequences were modeled independently, this choice limits the temporal dependencies available during training to those contained within each sequence.

Augmentation was applied only to the training split. The validation and test splits consisted exclusively of original observed sequences and were not synthetically resampled. For evaluation, these data were segmented into 10-s sequences, providing longer contiguous trial sequences for assessing model-generated behavior while reducing the frequency of sequence-boundary reinitialization. Across all three splits, sequences containing RT outliers or lacking at least one complete trial were excluded from subsequent analyses.

#### 4.4.3 Outlier removal

Trial sequences were excluded if any RT within the sequence was identified as an outlier. Outliers were defined using the median absolute deviation from the subject’s median (MAD), whereby trials whose RT absolute deviation exceeded 10× the MAD were excluded.

#### 4.4.4 Early stopping

To reduce overfitting, training was monitored using the validation split and terminated when performance no longer improved. We defined an early-stopping criterion, denoted by *J*_stop_, based on the discrepancy between model-derived and subject-derived switch costs on the validation data:

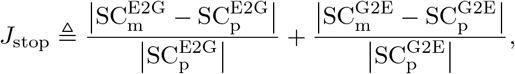

where 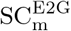 and 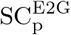 denote the model and subject switch costs for E2G trials, respectively, and 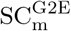 and 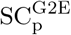 denote the corresponding quantities for G2E trials, respectively.

The stopping criterion *J*_stop_ was evaluated every 10 training epochs, beginning at epoch 500. Training was terminated if *J*_stop_ failed to decrease for 200 consecutive epochs. For all subsequent analyses, we retained the checkpoint corresponding to the minimum validation value of *J*_stop_. Model responses reported in the main analyses were always generated from the held-out test split, which was fully separate from the validation data used for early stopping. Unlike the RT summary statistics reported elsewhere, the switch costs used for early stopping were computed from all validation trials.

#### 4.4.5 Training hyperparameters

All GeoDynFormer models were trained using the same architecture and optimization hyperparameters, and all runs were initialized with the same random seed. Model training was performed on an NVIDIA RTX Ada 6000 GPU.

Optimization used the AMSGrad variant of the Adam algorithm ^(65)^, as implemented in PyTorch v1.9 ^(59)^. Unless otherwise noted, the optimizer hyperparameters were fixed at a learning rate of 10^−4^, *β*_1_ = 0.99, and *β*_2_ = 0.999. The training batch size was 2048.

To prevent instability from excessively large gradients, we applied gradient clipping with a threshold of 5. Specifically, when the Euclidean norm of the full parameter gradient exceeded 5, the gradient vector was rescaled to have norm 5. This norm was computed over all model parameters treated jointly as a single concatenated vector.

To compare model-generated behavior with observed behavior on the held-out test set, we computed trial-averaged RTs for each condition using only correct trials. Directional switch costs were then defined as *RT*_*E*2*G*_ − *RT*_*G*2*G*_ and *RT*_*G*2*E*_ − *RT*_*E*2*E*_, where each condition-level mean was computed from correct trials.

### 4.5 Latent-State and Trajectory Analysis

Unless otherwise specified, all analyses of model performance were benchmarked against observed behavior during the held-out test set. The Uniform Manifold Approximation and Projection (UMAP) ^(54)^ representation was used for visualization of latent trajectories and to motivate hypothesis-driven tests of latent state geometry. All statistical analyses of latent state geometry were conducted in the full 16-dimensional latent space except the orthogonality analysis performed on semantic axes derived from each subject’s UMAP embeddings.

#### 4.5.1 UMAP embedding of latent trajectories

To visualize the learned latent-state dynamics, as shown in Figs. 3a, 4, and 5, we projected the latent trajectories into a low-dimensional space using UMAP. For each fitted model, we collected the latent-state sequences from the held-out test set and arranged them into a latent-state matrix by concatenating all trajectories across time and trials. Specifically, if the latent trajectories had dimensions *T* × *N* × *D*, where *T* denotes the number of time steps, *N* is the number of trials, and *D* is the latent dimensionality, we reshaped them into a matrix of size (*T* × *N*) × *D*. UMAP was then fit to this matrix, and the resulting low-dimensional embedding was reshaped back into trajectory form for downstream analyses.

UMAP, implemented using the Python package `umap-learn` v0.5.3 ^(54)^, was configured with n components = 3, n neighbors = 100, min dist = 0.1, metric = cosine, random state = 1, and output metric = euclidean.

#### 4.5.2 Linear discriminant analysis

For the LDA analyses shown in Fig. 5b,c, latent trajectories from repeat trials were aligned to stimulus onset and grouped according to task type and gender×emotion attribute combinations. LDA models were trained using the latent state evaluated at the model’s mean RT, represented in the full 16-dimensional latent space. For each binary classification, the two classes were balanced by randomly downsampling the larger class to match the smaller class, and misclassification rates were estimated using 5-fold stratified cross-validation. The same cross-validation procedure was applied to the corresponding label-shuffled controls.

For the task type-classification analysis, LDA distinguished the gender and emotion task types while pooling across gender and emotion attribute combinations. For the general within-task classification analysis, each task type contained four gender×emotion attribute conditions: male–happy, male–angry, female–happy, and female–angry. Separate LDA models were trained for all six unique pairwise contrasts among these four conditions within each task type, and the resulting misclassification rates were averaged across the twelve comparisons to obtain a single summary value for each subject-specific model. For the task-relevant target attribute-classification analysis, classification of the correct task-relevant attribute was performed while holding the task-irrelevant attribute constant: within the gender task, male vs. female classification was performed separately for happy and angry attributes and then averaged, whereas within the emotion task, happy vs. angry classification was performed separately for male and female attributes and then averaged.

Shuffle controls were generated using 100 label permutations for each binary comparison, with each shuffled dataset evaluated using the same class-balancing and 5-fold stratified cross-validation procedure as the corresponding observed analysis. Following previous work ^(39)^, group-level statistical tests were performed using paired *t*-tests, comparing the observed misclassification rate with the mean of the corresponding null distribution. For the within-task classifications, shuffled misclassification rates were averaged across the constituent binary classifications within each shuffle iteration, followed by the same abovementioned procedure. Holm–Bonferroni correction was applied across the four LDA group-level comparisons: between-task classification, general within-task classification, gender-task task-relevant attribute classification, and emotion-task task-relevant attribute classification.

#### 4.5.3 Fixed-point estimation

Stable fixed points were identified by running each trained model in generative mode with long (50 s) sequences of static stimuli as inputs. When the latent dynamics converged to a stationary state, the converged latent configuration was treated as a stable fixed point. To determine whether convergence had occurred, we computed the standard deviation of each latent variable over the final 10 s of the sequence. If the mean of these standard deviations across latent dimensions was ≤0.001, the time-averaged latent state over the final 10 s was taken as a stable fixed point.

For each subject-specific model, fixed points were screened separately for all eight possible combinations of task type×gender attribute×emotion attribute. For each configuration, the model was initialized from 10 different random latent states sampled from latent positions visited across all time points and trials in the responses generated from the held-out test set. In nearly all cases, trajectories initialized from different starting states converged to the same endpoint for a given configuration. However, if the final latent state of a sequence was more than 0.001 in Euclidean distance from all other fixed points identified for that configuration, it was recorded as a distinct fixed point.

Stable fixed points associated with both task types were identified in all 22 subject-specific models. Across subjects, each model yielded exactly eight fixed points, corresponding to all combinations of task type×gender attribute×emotion attribute.

#### 4.5.4 Subject-specific semantic axes

Using the fixed points estimated for each repeat-trial condition, we constructed subject-specific semantic axes in the three-dimensional UMAP space (Fig. 3a; Extended Data Fig. 4). For each subject, fixed points were grouped by task type, gender attribute, and emotion attribute. From these condition-specific fixed-point locations, we defined three raw semantic contrast vectors corresponding to task type, task-relevant emotion, and task-relevant gender.

The task contrast vector was defined as the difference between the mean fixed-point locations for emotion-task and gender-task trials:

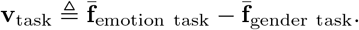

The task-relevant emotion contrast vector was defined within the emotion task as

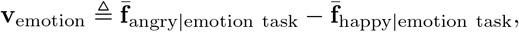

and the task-relevant gender contrast vector was defined within the gender task as

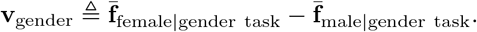

For the analysis of angular differences for target attributes across task relevance, the corresponding task-irrelevant contrasts were defined analogously, with emotion contrasted within the gender task and gender contrasted within the emotion task:

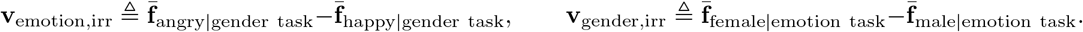

Separately, the three task-relevant raw contrast vectors were converted into an orthonormal subject-specific semantic basis for visualization. The task vector was first normalized to define **e**_task_. The emotion vector was then orthogonalized with respect to **e**_task_ and normalized to define **e**_emotion_. Finally, the gender vector was orthogonalized with respect to both **e**_task_ and **e**_emotion_ and normalized to define **e**_gender_. A cross-product fallback was implemented for numerical degeneracy, but no such degeneracy occurred in any of the subjects. This yielded the orthonormal subject-specific basis

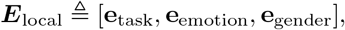

whose columns correspond to semantic axes expressed in the subject’s original UMAP coordinates.

To ensure consistent interpretation across subjects, sign conventions were enforced such that the positive directions corresponded to the emotion task on the task axis, angry expression on the emotion axis, and female gender on the gender axis. The resulting basis was aligned to a shared semantic reference frame with canonical axis order (task, emotion, gender), corresponding to [1, 0, 0], [0, 1, 0], and [0, 0, 1], respectively. The subject-specific rotation matrix was therefore

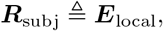

and any point **x** in the subject’s original UMAP space was projected into the common semantic reference frame as

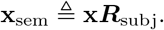

Pairwise angular relationships were assessed using the unit-normalized raw semantic contrast vectors 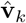 in the original three-dimensional UMAP space, prior to Gram–Schmidt orthogonalization ^(31)^. For the orthogonality analysis in Fig. 3b, the angular differences for task–gender, task–emotion, and emotion–gender were each tested against a prespecified direction of 90^◦^ using circular V-tests, with Holm– Bonferroni correction applied across these three comparisons. For the task-relevance analysis in Fig. 3c, the task-relevant and task-irrelevant emotion and gender contrast vectors were tested separately against a prespecified direction of 0^◦^; these two prespecified tests were not adjusted for multiple comparisons.

The omnibus orthogonality analysis likewise used the unit-normalized raw contrast vectors prior to orthogonalization. For the native-space control analysis, the same task, task-relevant emotion, and task-relevant gender contrasts were recomputed directly from the corresponding fixed points in the original 16-dimensional latent space. The group-level Gram matrix constructed in the original latent space showed non-zero off-diagonal values (task–emotion, −0.68; task–gender, −0.41; emotion–gender, 0.88). Consistently, the mean orthogonality error was substantially larger than expected for random triads of unit vectors in 16 dimensions (observed mean = 1.69; null mean = 0.57; 95% null interval (0.48, 0.66)).

#### 4.5.5 Latent-space distance measures

To quantify latent-space geometry, we performed three distance analyses: Euclidean distances between stable fixed points to characterize the hierarchical organization of the learned task representation (Fig. 5d,e); Euclidean distances between repeat- and switch-trial onset points to characterize onset point separation (Fig. 6a); and trial-wise Euclidean distances across consecutive states along each trajectory to assess the relationship between trajectory length and RT (Extended Data Fig. 6). All distance measures were computed directly in the full 16-dimensional latent space.

For the fixed-point analyses (Fig. 5d,e), Euclidean distances were computed between condition-specific stable fixed points. Between-task distances were calculated between G2G and E2E fixed points with face gender and emotion matched. Within-task distances were computed separately for task-relevant feature contrasts while matching the task-irrelevant feature, and for task-irrelevant feature contrasts while matching the task-relevant feature. For each contrast type, distances across all matched fixed-point pairs were averaged to obtain a single subject-level estimate.

For the switch-related onset-state analysis (Fig. 6a–e), all trajectories were aligned to stimulus onset, and an onset-state representative was defined for each trial condition as the centroid of the latent states evaluated at stimulus onset across all trials belonging to that condition. This yielded separate onset-state representatives for repeat and switch trials within each task type. For each switch direction, Euclidean onset point distance was computed between the switch-trial representative and the corresponding repeat-trial representative: E2G relative to G2G and G2E relative to E2E. Distances were first computed between switch and repeat conditions with matching face gender and emotion and then averaged across the four gender–emotion combinations separately for each switch direction. The same Euclidean onset point distance was used in the split-half reliability analysis (Fig. 6f,g).

For the trial-wise trajectory-length–RT analysis (Extended Data Fig. 6), cumulative latent trajectory length was computed separately for each trial from stimulus onset to that trial’s RT as the sum of Euclidean distances between consecutive latent states along the trajectory. Pearson’s *r* between trajectory length and RT was then computed across trials separately within each stimulus configuration, and the resulting correlation coefficients were averaged across configurations to obtain a single estimate for each subject-specific model.

A stimulus configuration was defined by the current trial’s task cue, face gender, and face emotion, yielding eight possible configurations. Configurations containing fewer than five trials were excluded from the averaging procedure. Two of 22 subject-level models had an excluded configuration, limited to one excluded configuration in each of the two models.

#### 4.5.6 Omnibus triad-level orthogonality analysis

To assess joint orthogonality among the task, task-relevant emotion, and task-relevant gender semantic axes, we analyzed the unit-normalized raw semantic contrast vectors prior to Gram–Schmidt orthogonalization. For each subject, the normalized raw semantic contrast matrix was defined as

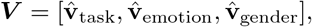

where 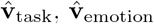, and 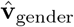 denote the unit-normalized raw task, task-relevant emotion, and task-relevant gender contrast vectors, respectively. The corresponding Gram matrix was computed as

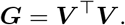

The diagonal entries of ***G*** equal 1, whereas the off-diagonal entries correspond to pairwise cosine similarities between the raw semantic contrast vectors (Fig. 3d). Under perfect orthogonality, ***G*** = ***I***, where ***I*** is the 3 × 3 identity matrix. Subject-level deviation from orthogonality was therefore quantified as

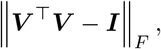

where ∥ · ∥_*F*_ denotes the Frobenius norm and smaller values indicate greater orthogonality.

To benchmark the observed group-mean orthogonality error against that expected for randomly oriented directions, we generated a dimension-matched Monte Carlo random-triad null distribution ^(18;17;32)^. For each iteration (*N* = 10, 000), one random triad was generated for each subject by drawing three independent Gaussian unit-normal vectors in the same ambient dimensionality as the observed semantic contrast vectors. The orthogonality error was computed using the same procedure, and then averaged across the matched number of subjects. A *p*-value was computed as the proportion of the null distribution less than or equal to the observed group mean.

#### 4.5.7 Split-half reliability analysis

For split-half reliability (internal consistency) analyses (Fig. 6f,g), the four mixed-task blocks were randomly partitioned across subjects into two non-overlapping halves.

Within each original block, behavioral sequences were segmented into 25-trial windows with a stride of a single trial; windows were not permitted to cross block boundaries. The number of retained windows was equalized between halves to the smaller available count. Within each half, windows were assigned to training, validation, and held-out test sets using the same 70%/10%/20% ratio as for the original GeoDynFormer models, and a separate GeoDynFormer was trained independently on each split half.

Within each independently trained half-model, raw switch cost and onset point distance were computed separately for E2G and G2E transitions using held-out test trials and the same procedure as for the subject-level models as described in Section 4.5.5.

Internal consistency was quantified across subjects as the Pearson correlation between the corresponding Half-1 and Half-2 estimates, separately for onset point distance and behavioral switch cost, each corrected using the Spearman–Brown formula. To compare the two internal consistency measures while accounting for uncertainty due to finite held-out trial sampling, we used a paired trial-level bootstrap with 5,000 iterations while keeping the subjects and independently fitted models fixed. Within each subject, split half, and switch direction, switch and corresponding repeat trials were resampled separately with replacement while preserving their original sample sizes. In each iteration, the corrected Pearson’s correlation was transformed using Fisher’s *r*-to-*z* transformation. Their difference was defined as

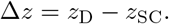

The bootstrap distribution was summarized by the fraction of iterations for which Δ*z >* 0–the proportion of bootstrapped samples in which onset point distance showed higher split-half reliability vs. the raw switch cost.

A two-sided paired-bootstrap *p*_boot_ value can be obtained by computing the proportion of bootstrapped differences in which Δ*z*_*b*_ ≤ 0 or Δ*z*_*b*_ ≥0, and taking the double of the smaller proportion. ^(14)^.

#### 4.5.8 Switch-trajectory transition point and tortuosity

For the switch-trajectory analyses shown in Fig. 7, we quantified epoch-specific trajectory geometry in the original 16-dimensional latent space. For each switch condition, we defined a transition point, *t*_*c*_, relative to the onset point of the repeat condition for the current task. Specifically, for E2G trials, *t*_*c*_ was defined as the time point along the E2G trajectory that minimized the Euclidean distance to the G2G onset point; for G2E trials, *t*_*c*_ was defined analogously relative to the E2E onset point. This partitioned each switch trajectory into an early epoch, [0, *t*_*c*_], and a late epoch, [*t*_*c*_, *RT*], where time 0 denotes stimulus onset and *RT* denotes the condition-specific mean reaction time.

For a trajectory segment spanning a generic interval [*t*_*a*_, *t*_*b*_], cumulative trajectory length was defined as

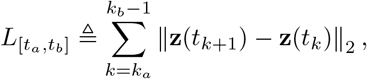

where **z**(*t*_*k*_) denotes the latent state at sampled time point *t*_*k*_ within the interval. The corresponding endpoint displacement was defined as the Euclidean distance between the first and last latent states of the same segment,

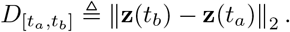

Trajectory tortuosity ^(33;6)^ was then defined as the ratio of cumulative path length to endpoint displacement,

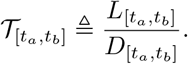

Thus, = 1 corresponds to a perfectly straight trajectory segment, whereas larger values indicate increasingly indirect trajectories. For each switch direction, we computed early trajectory length 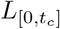, early trajectory tortuosity 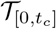, and late trajectory tortuosity 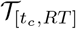 for subsequent analyses relating latent trajectory geometry to drift–diffusion model parameters and asymmetric switch costs.

#### 4.5.9 Trajectory alignment metrics

To quantify directional similarity between latent trajectories, we defined trajectory-alignment metrics based on the cosine similarity between condition-specific velocity vectors in the original 16-dimensional latent space. As illustrated in Fig. 8a,e, angular alignment was computed across condition pairs sharing either the same task-relevant attribute or the same task-irrelevant attribute (depending on the alignment metric) while matching the remaining attributes.

For a given pair of trajectories *A* and *B*, let **v**_*A*_(*t*) and **v**_*B*_(*t*) denote their velocity vectors at time *t*, estimated using numerical differentiation. The instantaneous alignment between the two trajectories was defined as

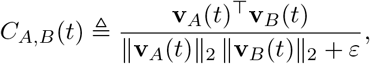

where *ε* is a small constant for numerical stability. Velocity vectors were evaluated on the discrete time samples of each extracted trajectory segment, referred to here as the trajectory’s time grid. When trajectories differed in duration or number of samples, as can occur for windows defined using condition-specific *t*_0_ or mean RT, each velocity trajectory was first mapped onto a shared phase axis spanning [0, 1]. Each velocity component was then interpolated using a piecewise cubic Hermite interpolating polynomial (PCHIP) ^(25;24)^ before cosine similarity was evaluated on a common phase grid.

For each condition pair, the cosine-similarity time course was summarized by its temporal mean over a specified interval [*t*_*a*_, *t*_*b*_],

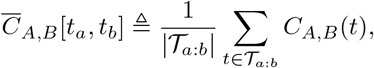

where *T*_*a*:*b*_ denotes the set of valid sampled time points within the interval [*t*_*a*_, *t*_*b*_]. For notational simplicity, the interval argument is omitted from the atomic condition-pair scores below when the comparison window is clear from context.

For each trial type *q* ∈ {G2G, E2E, E2G, G2E}, we computed four atomic condition-pair alignment scores:

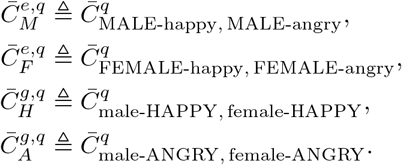

The first two scores compare trajectories with the same gender and different emotion, whereas the latter two compare trajectories with the same emotion and different gender. Thus, these scores quantify whether trajectories remain directionally aligned when the non-shared feature changes.

These atomic scores were then grouped into task-relevant and task-irrelevant trajectory-alignment metrics: the task-relevant alignment metric, *A*_rel_, was defined from trajectory pairs that shared the task-relevant attribute while differing in the task-irrelevant attribute; conversely, the task-irrelevant alignment metric, *A*_irr_, was defined from trajectory pairs that shared the task-irrelevant attribute while differing in the task-relevant attribute. For gender-task trials (G2G, E2G), this gives

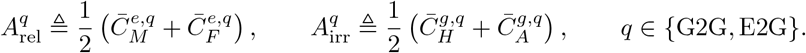

For emotion-task trials (E2E, G2E), the grouping was reversed:

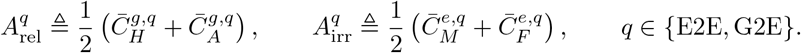

For the switch-trial analyses shown in Fig. 8a–d, *A*_irr_ was computed over the interval [0, *t*_0_], where time 0 denotes face onset. For the repeat-trial analyses shown in Fig. 8e–h, *A*_rel_ was computed over the interval [*t*_0_, *RT*], where *t*_0_ denotes the non-decision time estimated by the HDDM and *RT* denotes the condition-specific mean reaction time.

### 4.6 Drift–Diffusion Modeling

Hierarchical drift–diffusion modeling was used to estimate sub-process parameters from subjects’ RT distributions for each trial type (Fig. 7a). The hierarchical drift– diffusion model (HDDM) embeds the classical drift–diffusion model within a Bayesian hierarchical framework, allowing subject-specific parameters to be estimated while borrowing statistical strength from the group ^(76)^. We fit models using the HDDM Python toolbox ^(76)^. In this framework, observed RTs and choices are modeled as arising from latent decision-related sub-processes, including drift rate *v*, boundary separation *a*, starting bias *z*, and non-decision time *t*_0_. Because HDDM estimates posterior distributions over these parameters rather than single point estimates, it provides stable subject-level inference with fewer repeated trials per condition than non-hierarchical implementations (see Discussion).

In the present study, we adopted a simplified HDDM formulation to estimate the parameters most relevant for linking behavioral performance to the learned latent dynamics of GeoDynFormer. Specifically, we focused on drift rate *v*, reflecting the efficiency of evidence accumulation; boundary separation *a*, indexing response caution; and non-decision time *t*_0_, capturing processing stages outside the decision process itself, such as perceptual encoding, motor execution, and in the case of switch trials, task set reconfiguration. The resulting hierarchical Bayesian estimates provided subject-specific parameters that were directly related to the latent state trajectories learned by GeoDynFormer (Fig. 7; Extended Data Fig. 7).

Each task condition was fitted independently. To assess convergence, we fit three independent chains for each trial type using different random seeds. For repeat conditions, each chain was sampled for 15,000 iterations, with the first 3,000 samples discarded as burn-in. For switch conditions, each chain was sampled for 20,000 iterations, with the first 4,000 samples discarded as burn-in. Convergence was evaluated using the Gelman–Rubin diagnostic ^(27;11)^, and all fitted parameters were required to satisfy 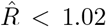. Only models meeting this criterion were retained for subsequent analyses.

### 4.7 Statistical Analysis

Unless otherwise specified, all reported statistical tests were two-sided, with statistical significance assessed at *α* = 0.05; for one-sided tests, statistical significance was assessed with corrected threshold *α* = 0.025. Unless otherwise noted, 95% confidence intervals were computed from *N* = 1,000 bootstrapped datasets. Where shown, shaded regression bands denote 95% bootstrapped confidence bands for fitted regression lines computed using the same resampling procedure. Additional analysis-specific details, including null-model and permutation procedures, are provided in the corresponding text. For boxplots, the center line indicates the median, the box limits indicate the upper and lower quartiles, whiskers extend to the most extreme data points within 1.5× the interquartile range, and points outside this range are shown as outliers.

For analyses involving repeated observations across conditions within subjects, we used repeated-measures correlation ^(5)^, as implemented in Pingouin v 0.5.3 ^(74)^, to estimate the common within-subject association while accounting for the non-independence of observations from the same subject. Holm–Bonferroni-adjusted *p*-values are denoted *p*_Holm_.

---

**Extended Data Fig 1:**
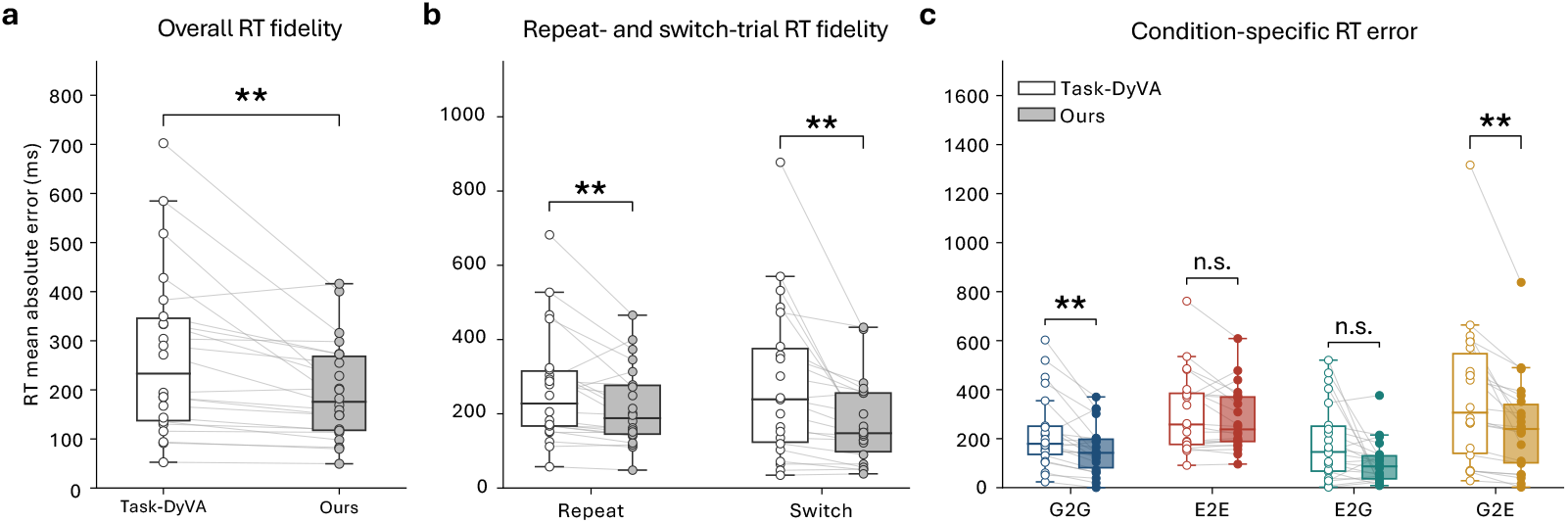
Transformer-based posterior inference improves recovery of condition-wise RTs relative to recurrent inference. The comparison model implemented the recurrent LSTM-based posterior-inference architecture of Jaffe et al. ^(39)^, whereas our model used Transformer-based posterior inference. Both models were trained on the same subject-level ATS datasets using the same ATS training framework and otherwise matched model-training settings, including the same data preprocessing, augmentation, optimization, and early-stopping procedure, thereby isolating the posterior-inference architecture. For each subject, the architecture comparison was evaluated using correct trials from the validation set at the checkpoint triggered by the early-stop criterion, *J*_stop_ in Sec. 4.4.4. The results below provide an empirical basis for using Transformer-based posterior inference in GeoDynFormer under an otherwise matched generative-model configuration. **a**, Mean absolute error (MAE) between model-generated and observed condition-wise mean RTs, averaged across all task conditions. RT MAE was lower with Transformer vs. recurrent inference (196.77 vs. 272.47 ms; paired *t*(21) = 3.34, *p* = 0.0031). Note that although we refer to the first model as Task-DyVA, it was identical to GeoDynFormer except for the inference model. **b**, RT MAE stratified across repeat and switch conditions. Transformer-based inference yielded lower RT error for both repeat (217.6 vs. 265.7 ms; paired *t*(21) = 2.86, *p*_Holm_ = 0.0094) and switch trials (175.9 vs. 279.2 ms; paired *t*(21) = 3.28, *p*_Holm_ = 0.0072); correction was applied across these two conditions. **c**, RT MAE shown separately for each condition (G2G, E2E, E2G, G2E). Transformer-based inference yielded lower error for G2G (156.1 vs. 225.6 ms; paired *t*(21) = 3.58, *p*_Holm_ = 0.0053) and G2E (250.9 vs. 372.6 ms; paired *t*(21) = 4.06, *p*_Holm_ = 0.0023) trials, whereas differences were not significant for E2E (279.1 vs. 305.8 ms; paired *t*(21) = 1.72, *p*_Holm_ = 0.101) and E2G (101.0 vs. 185.8 ms; paired *t*(21) = 2.40, *p*_Holm_ = 0.052) trials; correction was applied across the four comparisons. For all panels, *N* = 22 paired subject-level models; points indicate individual subjects and lines connect corresponding recurrent and Transformer models. Open and filled markers indicate recurrent and Transformer-based inference, respectively. All statistical comparisons were paired *t*-tests. ∗∗ *p <* 0.01, otherwise n.s.

**Extended Data Fig 2:**
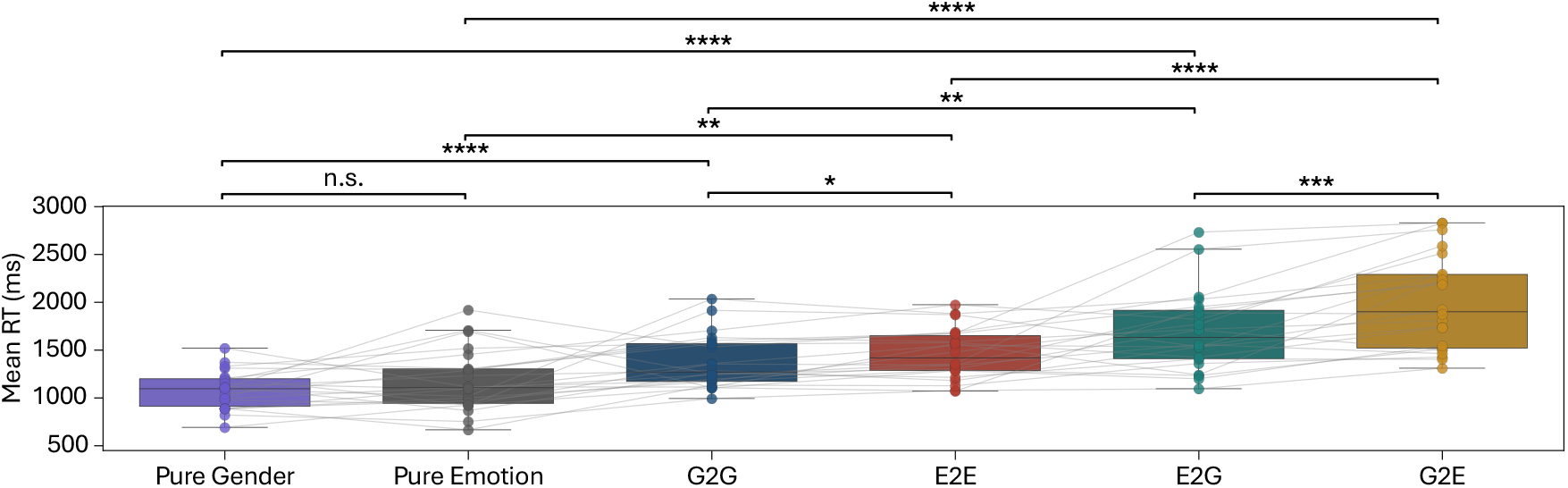
Mean RTs across pure, repeat, and switch trial types. Group mean ± s.e.m. was 1079 ± 43 ms for pure gender (from single-task blocks), 1174 ±69 ms for pure emotion, 1375 ±59 ms for G2G, 1457 ±56 ms for E2E, 1692 ±89 ms for E2G, and 1996 ±106 ms for G2E. Paired *t*-test results: pure gender vs. pure emotion, *t*(21) = −1.48, n.s.; G2G vs. E2E, *t*(21) = −2.71, *p* = 0.013; E2G vs. G2E, *t*(21) = −4.48, *p <* 10^−3^; pure gender vs. G2G, *t*(21) = −5.35, *p <* 10^−4^; pure emotion vs. E2E, *t*(21) = −3.72, *p <* 0.01; G2G vs. E2G, *t*(21) = −3.79, *p <* 0.01; E2E vs. G2E, *t*(21) = −6.22, *p <* 10^−5^; pure gender vs. E2G, *t*(21) = −6.30, *p <* 10^−7^, pure emotion vs. G2E, *t*(21) = −8.71, *p <* 10^−7^. Points represent individual subjects, boxplots show the median and 25th–75th percentiles, whiskers extend to the most extreme values within 1.5× the interquartile range, and gray lines connect paired observations. ∗ *p <* .05, ∗∗ *p <* .01, ∗∗∗ *p <* .001, ∗∗∗∗ *p <* .0001, otherwise n.s.

**Extended Data Fig 3:**
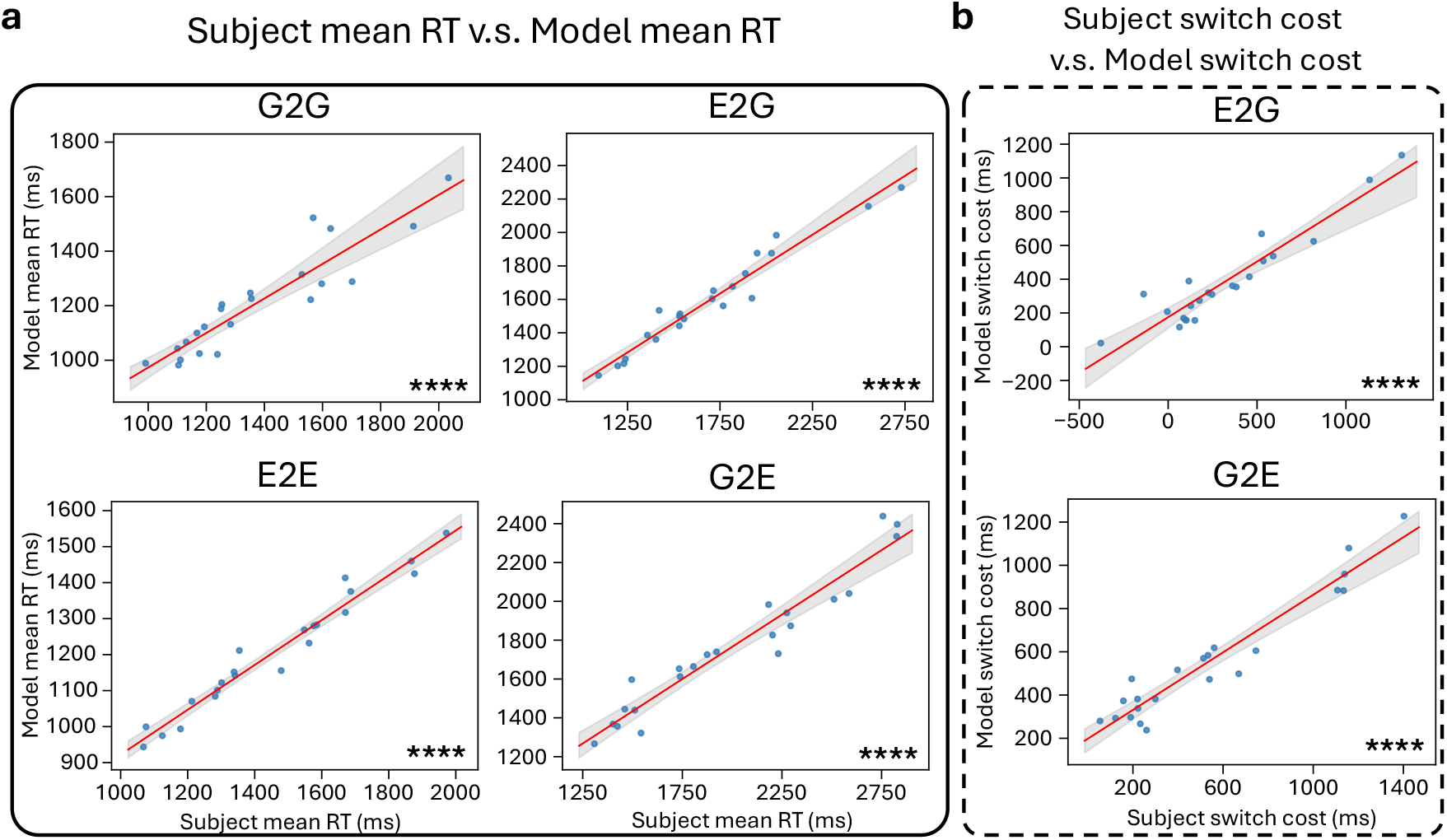
Observed subject-level data vs. subject-level model correspondence in condition-wise mean RTs and switch costs. **a**, Observed vs. model-generated mean RTs, shown separately for G2G (Pearson’s *r* = 0.92, bootstrap 95% CI (0.84, 0.97), *p <* 10^−8^), E2E (Pearson’s *r* = 0.98, bootstrap 95% CI (0.96, 0.99), *p <* 10^−14^), E2G (Pearson’s *r* = 0.98, bootstrap 95% CI (0.94, 0.99), *p <* 10^−13^), and G2E (Pearson’s *r* = 0.96, bootstrap 95% CI (0.93, 0.99), *p <* 10^−12^). **b**, Observed vs. model-generated switch cost for E2G (top; Pearson’s *r* = 0.94, bootstrap 95% CI (0.82, 0.98), *p <* 10^−10^) and G2E (bottom; Pearson’s *r* = 0.96, bootstrap 95% CI (0.91, 0.98), *p <* 10^−11^). For **a** and **b**, points represent one subject–model pair; red lines indicate best linear fits, and gray shaded regions 95% bootstrapped CIs, *N* = 22. ∗∗∗∗ *p <* .0001.

**Extended Data Fig 4:**
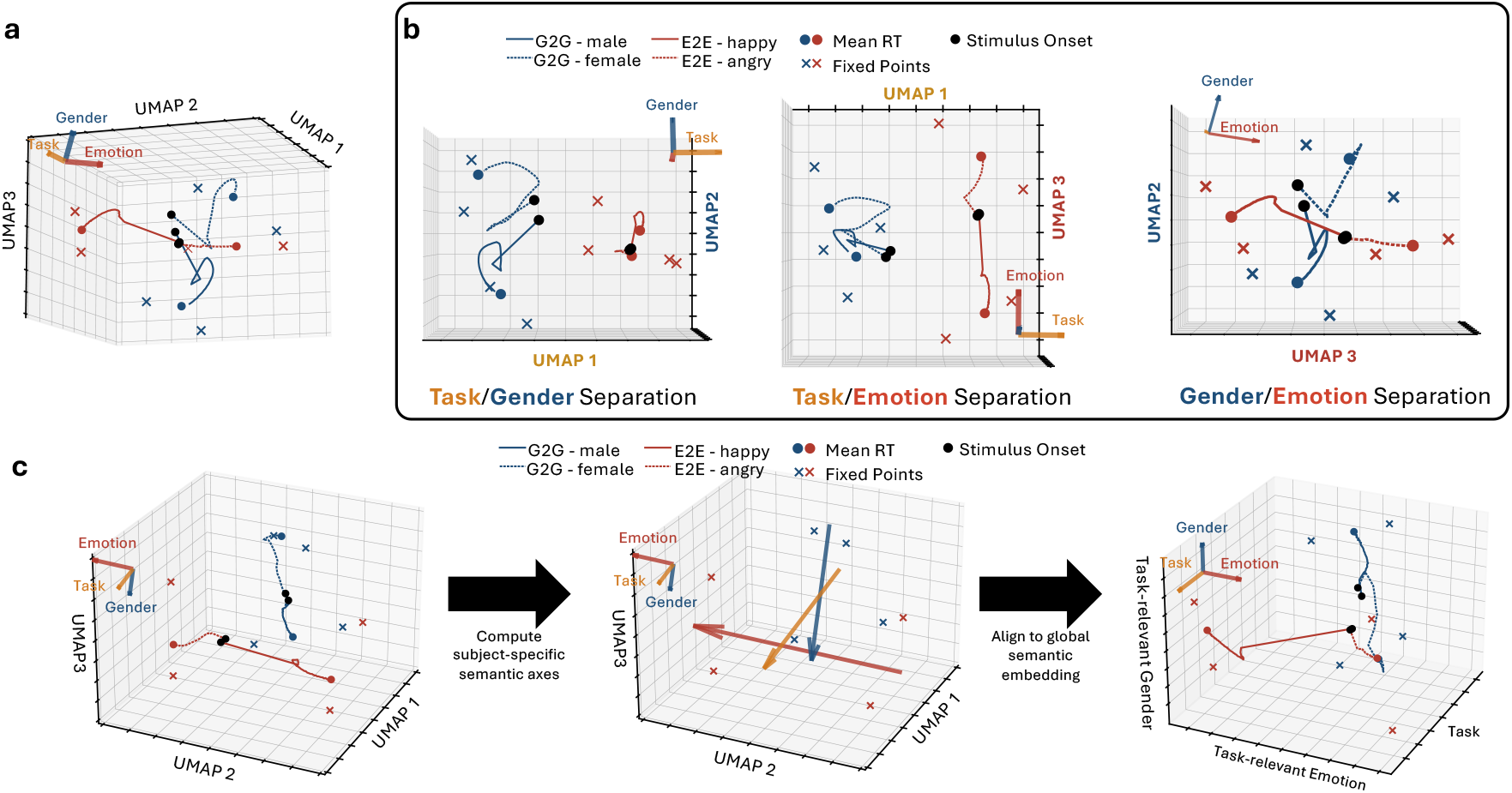
Additional examples of subject-level UMAP embeddings of repeat-trial latent trajectories and alignment to shared semantic embedding. **a**, Expanded visualization for the same example subject shown in Fig. 3a. Trial-averaged repeat-trial trajectories in the original 3D UMAP embedding with stimulus-onset points, mean-RT points, and fixed points. **b**, Complementary 2D UMAP projections showing latent trajectories stratified across task type×task-relevant gender attribute, task type×task-relevant emotion attribute, and task-relevant gender attribute×task-relevant emotion attribute. **c**, Additional single-subject example of semantic-axis construction and alignment to shared semantic embedding. Left, repeat-trial trajectories in the subject’s original 3D UMAP embedding. Middle, semantic axes computed from fixed points. Right, the same trajectories projected into the common semantic embedding with task type, task-relevant emotion, and task-relevant gender axes.

**Extended Data Fig 5:**
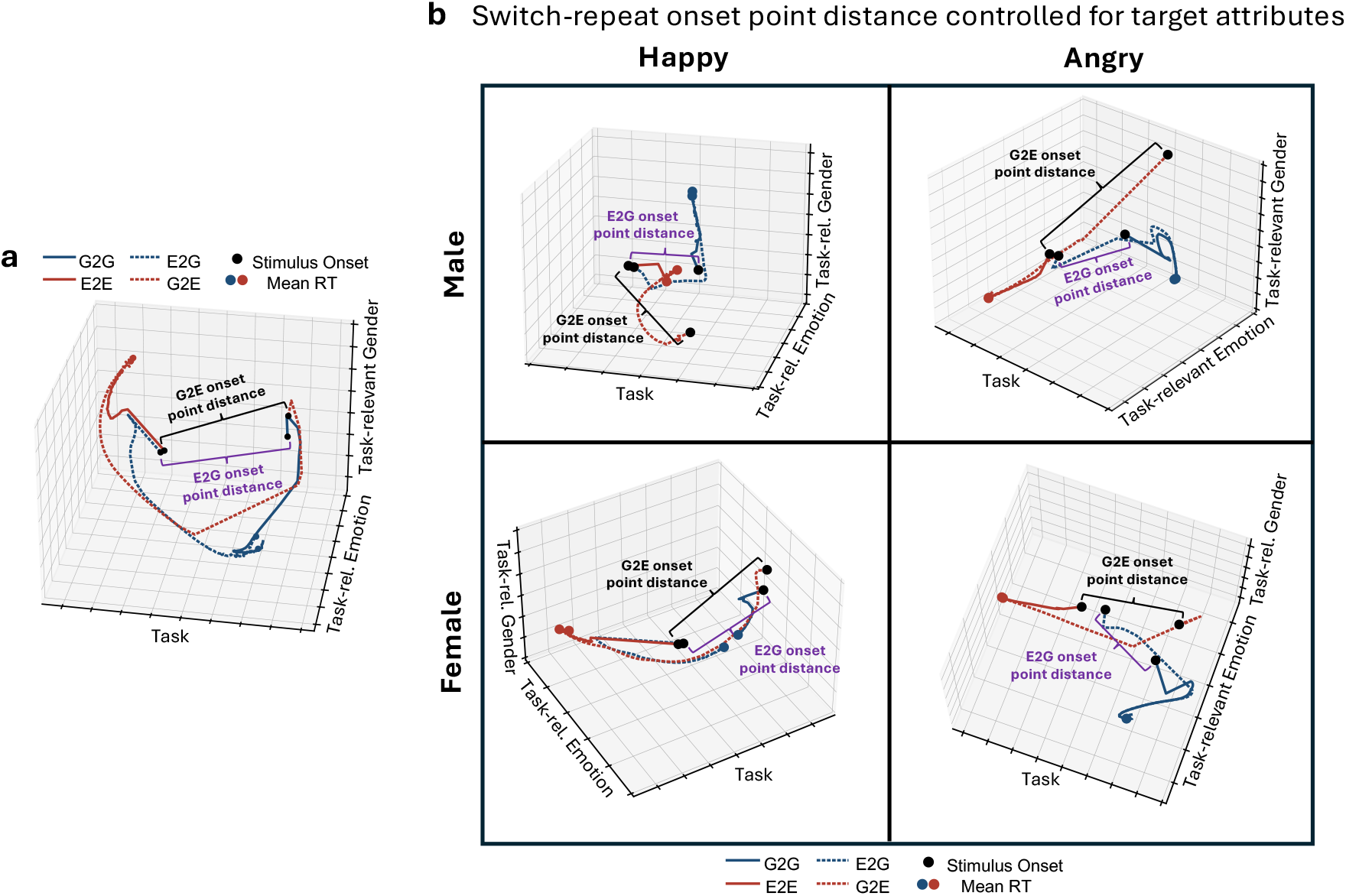
Organization of switch and repeat trials in shared semantic embedding and quantification of onset point separation controlling for target attribute conditions. **a**, Trial-averaged switch-trial trajectories in common semantic embedding. Single-subject example illustrates the organization of E2G and G2E trajectories relative to stimulus-onset and mean-RT points. **b**, Conceptual illustration of the onset point distance metric in shared semantic embedding, which controlled for the four target attribute combinations. For each of these target attribute conditions, the distance was computed between the switch-trial onset point and the corresponding repeat-trial onset point. The reported onset point distance metric took the average across these four target attribute conditions; onset point distance was computed in the original 16D latent space for downstream statistical analyses.

**Extended Data Fig 6:**
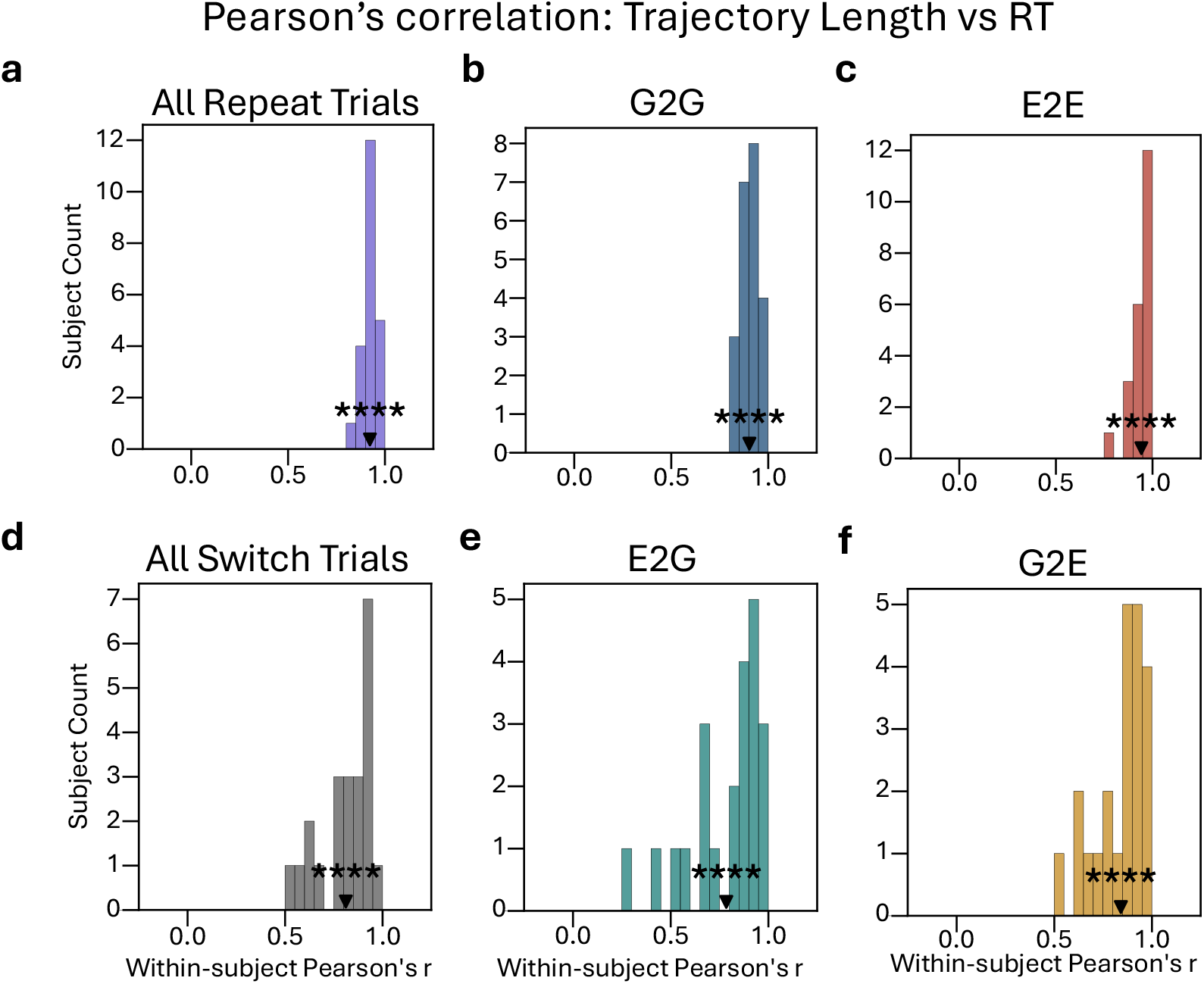
Within-subject correlation between cumulative latent trajectory length and RT across trials. For each subject-specific model, Pearson’s *r* between cumulative trajectory length and RT was computed across trials separately within each stimulus configuration and then averaged across configurations to obtain a subject-level correlation coefficient. For all panels, black triangles indicate the group-level *r*. One-sample *t*-tests against zero gave the following results: all repeat trials (mean ± s.e.m. = 0.92 ± 0.0086, *t*(21) = 107.50, *p <* 10^−29^), G2G (0.90 ± 0.0097, *t*(21) = 93.23, *p <* 10^−28^), E2E (0.94 ± 0.010, *t*(21) = 93.22, *p <* 10^−28^), all switch trials (0.81 ± 0.028, *t*(21) = 28.80, *p <* 10^−17^), E2G (0.78 ± 0.041, *t*(21) = 19.09, *p <* 10^−14^), and G2E (0.84 ± 0.027, *t*(21) = 30.91, *p <* 10^−18^). ∗∗∗∗ *p <* .0001.

**Extended Data Fig 7:**
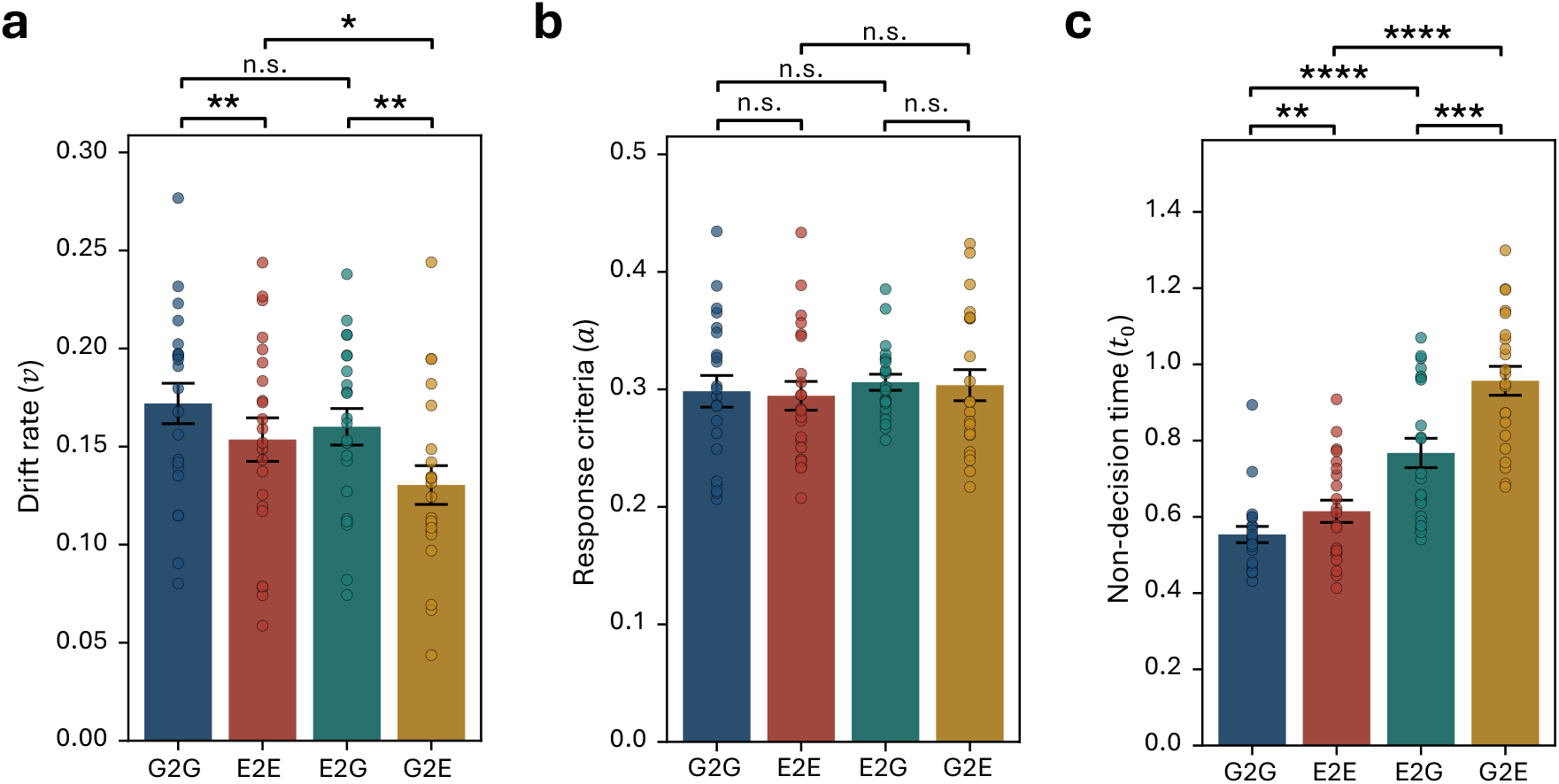
DDM parameter estimates using HDDM ^(76)^ across ATS conditions. **a**, Drift rate (*v*) across G2G, E2E, E2G, and G2E. Paired *t*-test results: G2G vs. E2E, *t*(21) = 3.48, *p*_Holm_ = 0.0090; E2G vs. G2E, *t*(21) = 4.42, *p*_Holm_ = 0.0012; G2G vs. E2G, *t*(21) = 1.54, *p*_Holm_ = 0.28; E2E vs. G2E, *t*(21) = 2.92, *p*_Holm_ = 0.024. **b**, Boundary separation (*a*); no pairwise differences were significant after Holm–Bonferroni correction (all adjusted *p* = 1.00). **c**, Non-decision time (*t*_0_). G2G vs. E2E, *t*(21) = −3.13, *p*_Holm_ = 0.0050; E2G vs. G2E, *t*(21) = −5.09, *p*_Holm_ *<* 10^−3^; G2G vs. E2G, *t*(21) = −6.29, *p*_Holm_ *<* 10^−4^; E2E vs. G2E, *t*(21) = −9.08, *p*_Holm_ *<* 10^−7^. Holm–Bonferroni correction was applied across all six pairwise comparisons separately for each DDM parameter. Bars indicate group means, error bars indicate s.e.m., points indicate individual subjects, and brackets indicate paired *t*-test comparisons. ∗ *p <* .05, ∗∗ *p <* .01, ∗∗∗ *p <* .001, ∗∗∗∗ *p <* .0001, otherwise n.s.

**Extended Data Fig 8:**
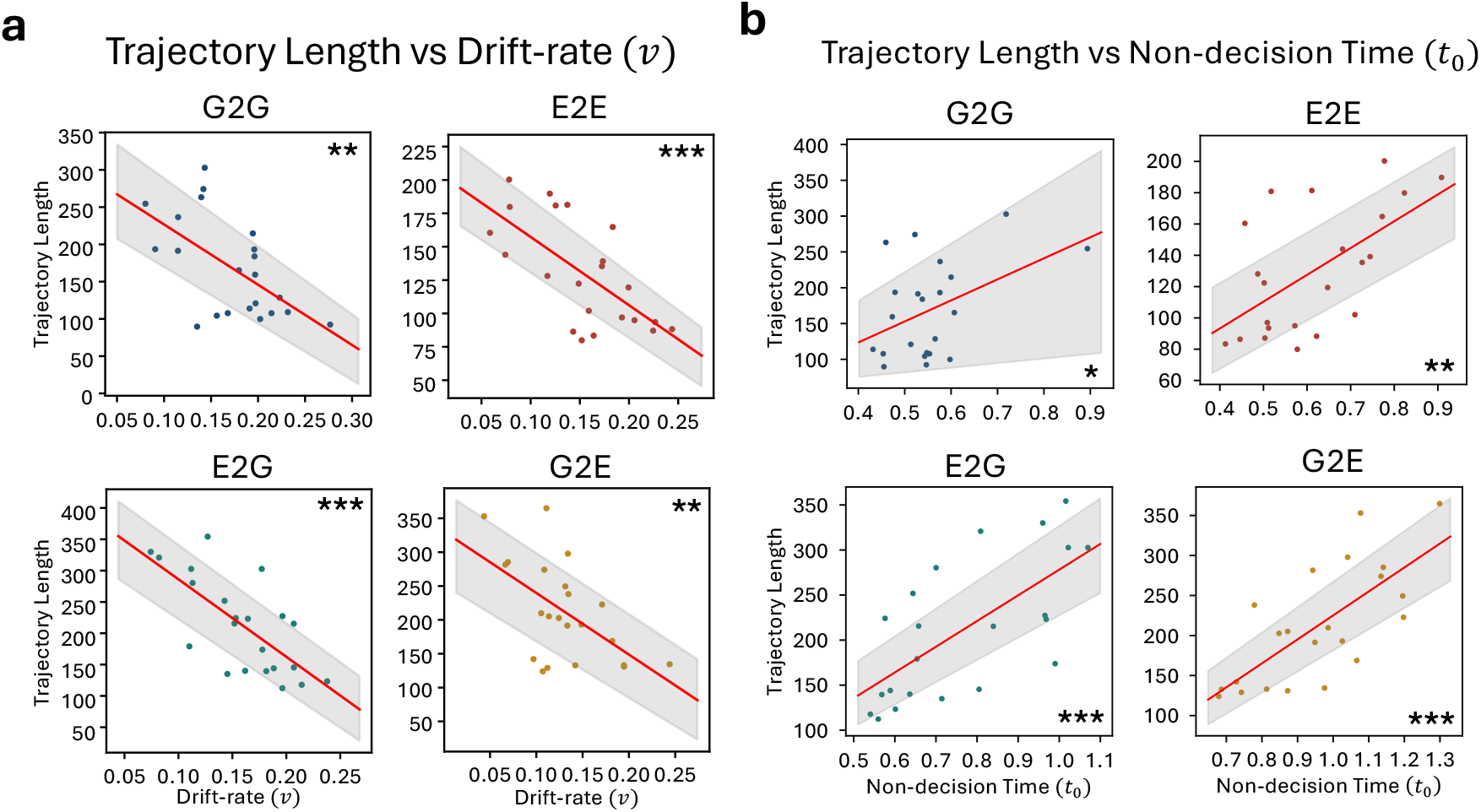
Latent trajectory length from the stimulus-onset point to the mean-RT point is associated with drift rate and non-decision time across ATS conditions. **a**, Cross-subject correlation between trajectory length (from the stimulus-onset point to the mean-RT point) vs. drift rate (*v*), shown separately for G2G (top left; Pearson’s *r* = −0.59, 95% CI (−0.79, −0.33), *p <* 0.01), E2E (top right; Pearson’s *r* = −0.67, 95% CI (−0.82, −0.46), *p <* 10^−3^), E2G (bottom left; Pearson’s *r* = −0.70, 95% CI (−0.88, −0.41), *p <* 10^−3^), and G2E (bottom right; Pearson’s *r* = −0.57, 95% CI (−0.81, −0.25), *p <* 0.01) conditions. **b**, Trajectory length vs. non-decision time (*t*_0_), shown separately for G2G (top left; Pearson’s *r* = 0.44, 95% CI (− 0.09, 0.73), *p* = 0.040), E2E (top right; Pearson’s *r* = 0.59, 95% CI (0.23, 0.84), *p* = 0.0040), E2G (bottom left; Pearson’s *r* = 0.67, 95% CI (0.40, 0.87), *p <* 10^−3^), and G2E (bottom right; Pearson’s *r* = 0.72, 95% CI (0.48, 0.87), *p <* 10^−3^) conditions. For all panels, points represent one subject–model pair (*N* = 22); red lines indicate best linear fits; gray shaded regions indicate 95% bootstrapped CI. ∗*p <* .05, ∗∗ *p <* .01, ∗∗∗ *p <* .001.

**Extended Data Fig 9:**
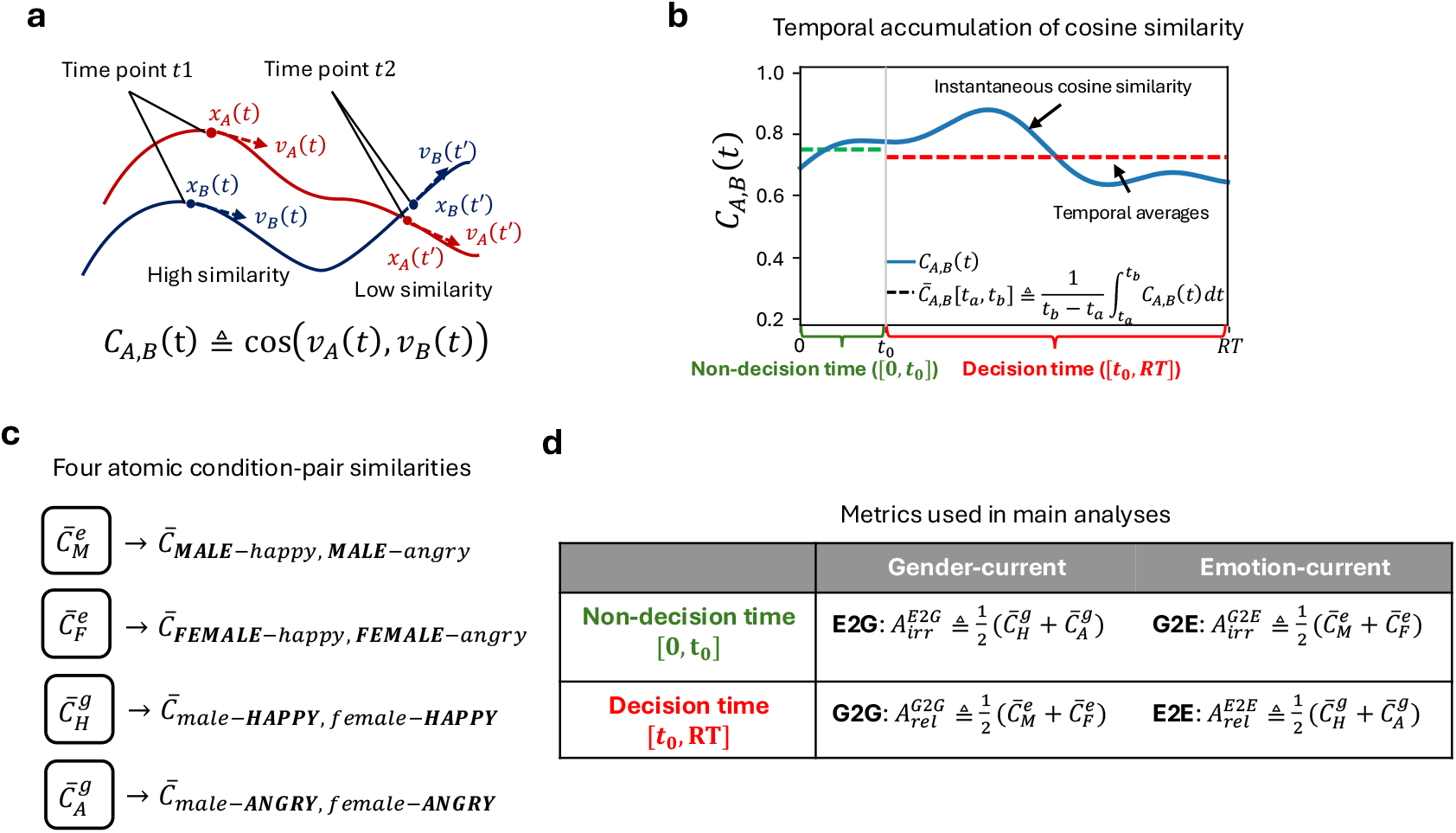
Definition of angular alignment metrics across trajectory pairs. **a**, Instantaneous cosine similarity between two latent trajectories, computed from their velocity vectors at each time point. **b**, Temporal averaging of instantaneous cosine similarity over a specified interval [*t*_*a*_, *t*_*b*_] produced a condition-pair alignment score, 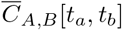. The early non-decision time epoch [0, *t*_0_] and late decision time epoch [*t*_0_, *RT*] used in the main analyses are indicated. **c**, Four atomic angular similarities represent the time-averaged cosine similarity across the two emotion attributes while holding gender attributes fixed (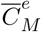 and 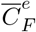), or across the two gender attributes while holding emotion attributes fixed (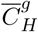 and 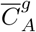). **d**, The flexible task-set theory ^(71)^ posits that the gender task set is more strongly shielded and shows greater cross-trial carry-over (inertia) than the emotion task set. To test this theory, we defined task-relevant (*A*_rel_) and task-irrelevant (*A*_irr_) angular alignment metrics that quantified the angular similarity of latent trajectories across conditions sharing the same task-relevant or task-irrelevant target attribute, respectively. During non-decision time ([0, *t*_0_]), *A*_irr_ was used to assess switch trajectories: for E2G, 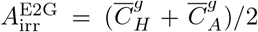, whereas for G2E, 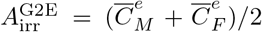. During decision time ([*t*_0_, *RT*]), *A*_rel_ was used to assess repeat trajectories: for G2G, 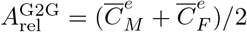, whereas for E2E, 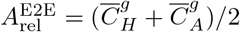.

